# TTBK2 controls cilium stability through actin and the centrosomal compartment

**DOI:** 10.1101/2022.06.02.494590

**Authors:** Abraham Nguyen, Sarah C. Goetz

## Abstract

The serine-threonine kinase Tau Tubulin Kinase 2 (TTBK2) is a key regulator of the assembly of primary cilia, which are vital signaling organelles. TTBK2 is also implicated in the stability of the assembled cilium, through mechanisms that remain to be defined. Here, we use mouse embryonic fibroblasts (MEFs) derived from *Ttbk2^fl/fl^; UBC-CreERT+* embryos (hereafter *Ttbk2^cmut^*) to dissect the role of TTBK2 in cilium stability. This system depletes TTBK2 levels after cilia formation, allowing us to assess the molecular changes to the assembled cilium over time. As a consequence of *Ttbk2* deletion, the ciliary axoneme is destabilized and primary cilia are lost within 48-72 hours following recombination. Axoneme destabilization involves an increased frequency of cilia breaks and is partially driven by altered actin dynamics and a reduction in axonemal microtubule modifications. At the same time, we find that TTBK2 is required to regulate the composition of the centriolar satellites and to maintain the basal body pools of intraflagellar transport (IFT) proteins. Altogether, our results reveal parallel pathways by which TTBK2 maintains cilium stability.

## Introduction

Primary cilia are critical regulators of a variety of different cell signaling pathways, including Sonic Hedgehog (SHH) and G-protein coupled receptor (GPCR) signaling (1,2). While most vertebrate cell types have a primary cilium, these structures are dynamic and are assembled and disassembled during development in response to cell cycle as well as extracellular cues (3). While there has been progress in uncovering key players and molecular steps involved in primary cilium assembly, less is known about the regulation of cilium disassembly, or the molecular players and pathways governing stability and maintenance of the assembled ciliary axoneme. Defining the regulation of cilia at each distinct phase of the cilia lifecycle is important for our understanding of how critical signal transduction pathways are regulated.

Cilium assembly is initiated at the mother centriole, the older of the cell’s two centrioles, which is distinguished from the daughter centriole by the presence of sub-distal (SDA) and distal appendages (DA). These appendages are important for organizing microtubules, and for recruiting the machinery that will assemble the cilium, respectively (5). DA proteins including CEP164 are sequentially recruited to the mother centriole (6) downstream of the distal centriolar protein oral-facial-digital-1 (OFD1) (7), and in turn mediate the recruitment of the ciliary kinase TTBK2 as well as ciliary vesicles that will ultimately give rise to the ciliary membrane (6,8). Intraflagellar transport (IFT) proteins are then recruited to assemble the cilium by trafficking tubulin and other proteins to assemble the ciliary axoneme which will project from the cell surface (8,9).

In previous studies, we identified a requirement for TTBK2 in the initial stages of cilium assembly, showing that TTBK2 is required both to recruit IFT proteins to the basal body and to remove suppressors of cilium assembly such as CP110 (10). Thus, in cultured cells and mouse embryos lacking TTBK2, cilium assembly fails, resulting in mid-gestation lethality (10). Using a mouse allelic series for *Ttbk2*, we subsequently demonstrated that this kinase is also important for the structure and stability of the ciliary axoneme, and for ciliary trafficking (11). Furthermore, deletion of *Ttbk2* in adult mice results in reduced cilia frequency, suggesting that TTBK2 is required to maintain the structure of cilia as well as to initiate its assembly (12). However, the cellular and molecular mechanisms by which TTBK2 regulates both cilium initiation and maintenance remain largely unknown.

In this work, we inducibly delete *Ttbk2* from cells in culture to probe its roles in cilium stability. We show that *Ttbk2* deletion leads to progressive cilium instability and an increased frequency of primary cilia fragmentation leading to cilia loss. This cilia loss is partially attenuated by small molecule inhibitors affecting actin dynamics. Furthermore, *Ttbk2* deletion results in dysregulation of distinct elements of the centrosomal compartment at the base of the primary cilium including the loss of IFT pools and changes in the centriolar satellite composition. This study reveals novel roles for TTBK2 in the maintenance of primary cilia integrity and new insights into requirements for primary cilia stability.

## Results

### Inducible genetic removal of TTBK2 results in ciliary breakage and loss

TTBK2 is recruited to the basal body by DA proteins, where it is essential for early steps of cilium initiation. TTBK2 then remains at the basal body following the completion of ciliary axoneme assembly, but is lost from this compartment upon re-addition of serum to induce ciliary disassembly (10). Deletion of *Ttbk2* from adult tissues results in loss of cilia (12), implying that TTBK2 might be continuously required at the basal body to promote ciliary trafficking and/or stability. Our analysis of *Ttbk2* hypomorphic mutants identified structural abnormalities in the cilia of cells with reduced levels of TTBK2 protein (11).

To investigate the cellular requirements for TTBK2 in regulating primary cilia stability, we employed MEFs derived from *Ttbk2^fl/fl^* embryos also expressing a tamoxifen (Tmx)-inducible Cre recombinase under the control of the ubiquitin C promoter (*UBC-CreERT2+*) to allow for genetic removal of *Ttbk2* under temporal control. In the absence of Tmx, these cells, like other fibroblasts, form cilia when cultured in low-serum (0.5% FBS) conditions, and the percentage of ciliated cells is indistinguishable from WT MEFs (**Fig. S1A**). *Ttbk2^cmut^*MEFs were then cultured in low-serum media for 15 hours to induce cilia formation and treated with Tmx (1µM) to induce Cre-mediated recombination or with the vehicle (Veh), ethanol, as a control. T0 is defined as the point at which Tmx was administered. Tmx and Veh-treated cells were imaged 0, 24, 48, and 72 hours following Tmx treatment (**Fig. 1A**). We found that TTBK2 protein was reduced at the mother centriole within 24-48 hours post Tmx treatment (**Fig. 1B**), as is *Ttbk2* mRNA measured by qPCR (**Fig. S1B**). Cilia frequency was comparable in the Tmx-treated cells relative to Veh-treated for the first 24 hours (66.244 ± 4.705% vs 72.891 ± 5.258 %). By 48 hours, cilia frequency in Tmx-treated cells was less than half that of Veh-treated cells (29.66% ± 2.662% vs 70.00% ± 3.46%) and cilia were nearly gone (10.989 ± 2.549% vs 71.299 ± 3.213%) in the Tmx-treated cells 72 hours following treatment with Tmx (**Fig. 1C**). Treatment of *Ttbk2^fl/fl^,UBC-CreERT2-* cells with Tmx did not cause loss of cilia, confirming that the cilia loss was not an artifact of Tmx treatment (**Fig. S1C**). Stable expression of TTBK2-GFP in Tmx-treated *Ttbk2^cmut^*MEFs rescued cilia loss, confirming that this is due to specific loss of TTBK2 (**Fig. S1D**). Thus, this system recapitulates the requirements we have previously identified for TTBK2 in initiating cilium assembly (10) and stability (11).

**Figure 1:**
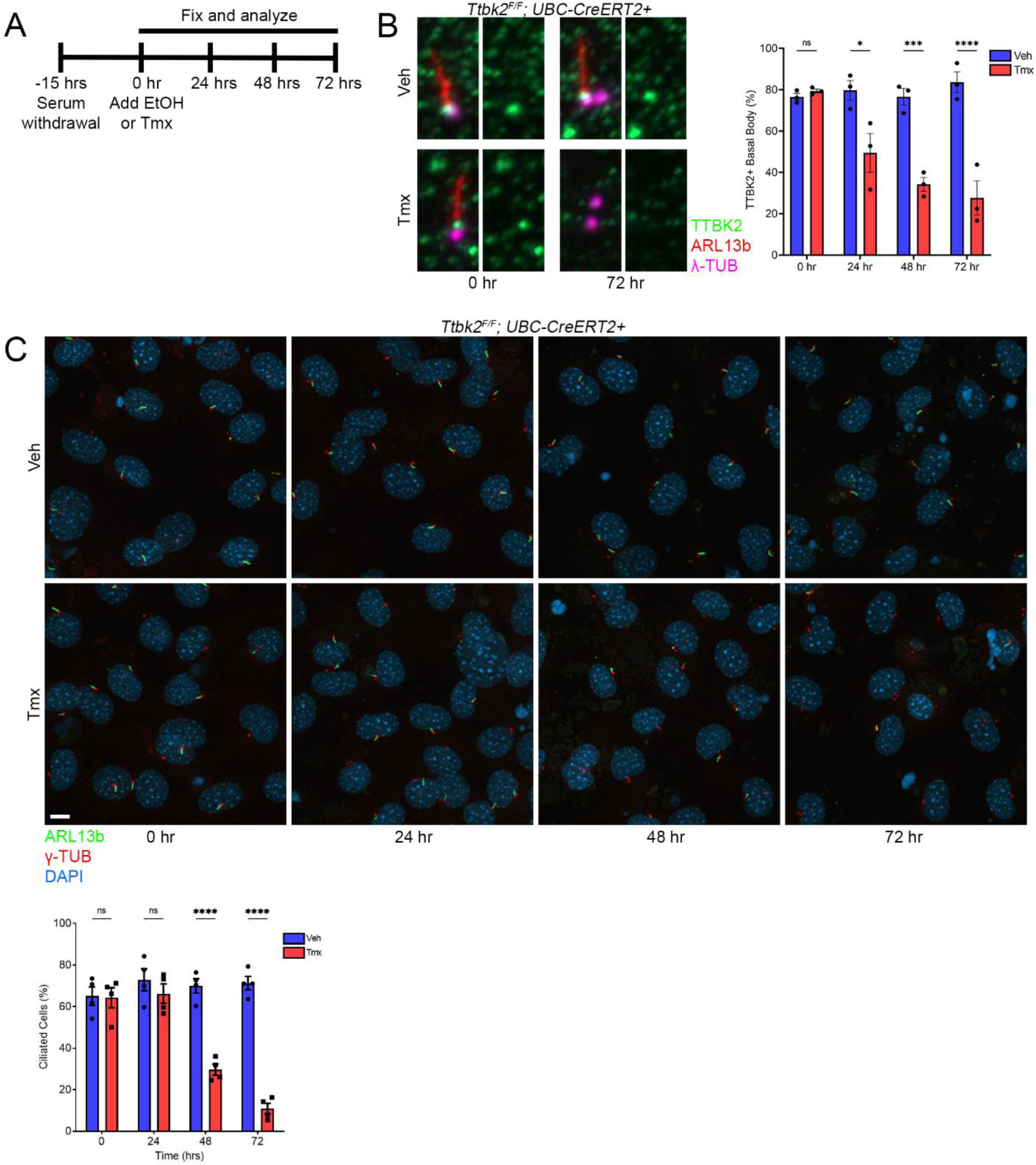
TTBK2 is required for maintaining primary cilia. A. Schematic of experimental design. *Ttbk2fl/fl; UBC-CreERT2+* MEFs were serum starved for 15 hours before being treated with ethanol (Veh) or tamoxifen (Tmx). MEFs were then fixed at the indicated time points. B. Representative images of *Ttbk2fl/fl; UBC-CreERT2+* MEFs treated with Veh or Tmx for 0 and 72 hours and immunostained for TTBK2 (green), ARL13b (red), and γ-Tubulin (magenta). Graph depicts mean percentage of basal bodies with positive TTBK2 staining in *Ttbk2fl/fl; UBC-CreERT2+* MEFs after treatment with Veh or Tmx for the indicated durations. Each dot represents an experiment with >50 cells. Statistical comparison was performed by two-way ANOVA with Tukey’s multiple comparisons test, * denotes p<0.05, *** denotes p<0.001, **** denotes p<0.0001. C. Representative images of *Ttbk2fl/fl; UBC-CreERT2+* MEFs treated with Veh or Tmx for 0, 24, 48, and 72 hours and stained for ARL13b (green), γ-Tubulin (red), and DAPI (blue). Scale bar: 10μm. Graph depicts mean percentage of ciliated cells ± SEM. Each dot represents an experiment with >50 cells. Statistical comparison was performed by two-way ANOVA with Tukey’s multiple comparisons test, **** denotes p<0.0001.

Having established an inducible system to specifically interrogate the cellular and molecular role of TTBK2 in ciliary stability, we next used this system for live-cell imaging. To follow cilia in real-time in the *Ttbk2^cmut^* MEFs ± Tmx, we crossed *Ttbk2^cmut^* mice to a double transgenic line expressing *Arl13b-mCherry* and *Centrin2-GFP* (*ACCG* mice)(13-15), and derived MEFs from *Ttbk2^cmut^;ACCG* embryos. We verified that loss of cilia in *Ttbk2^cmut^;ACCG* MEFs treated with Tmx takes place in the same overall timeframe as described for *Ttbk2^cmut^* cells (**Fig. S1E**). Given the sharp change in the frequency of ciliated cells from 24-48 hours and from 48-72 hours, we performed live cell imaging in a 20-hour window within these critical periods, 30-50 hours following treatment of the cells with Veh or Tmx (**Supplemental Videos 1-3**). Analysis of these live imaging studies revealed that the cilia of *Ttbk2^cmut^;ACCG* cells treated with Tmx underwent frequent breakages (**Fig. 2A**). A small percentage of vehicle-treated cells (5.44%) exhibited breakages from the distal tip of the cilium (referred to as “budding”), whereas about a third of Tmx-treated *Ttbk2^cmut^;ACCG* cells (29.18%) had one or more such budding events (**Fig. 2B**).

**Figure 2:**
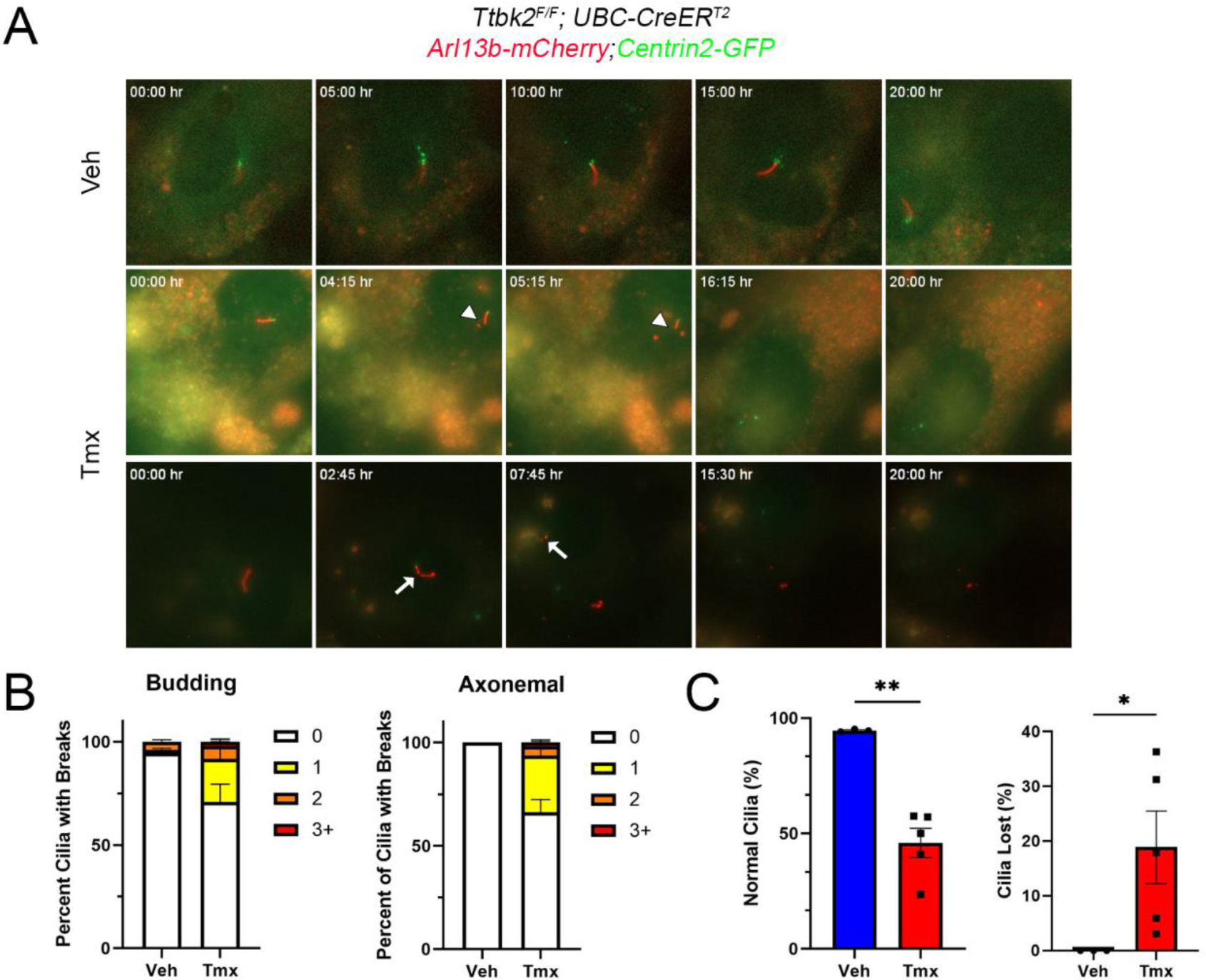
TTBK2 loss results in increased cilia fragmentation leading to cilia loss. A. *Ttbk2^cmut^;ACCG* MEFs were serum starved for 15 hrs before being treated with Veh or Tmx. Live imaging was then performed from 30 hours to 50 hours after induction. Arrowheads show examples of cilia breakages at the tips, termed “budding”. Arrows show examples of cilia breakages along the axoneme, termed “axonemal”. Time of video denotes hours:minutes into recording. B. Graph depicts mean percentage of cilia observed having one, two, or ≥ three budding events or axonemal breaks from three and five independent experiments where *Ttbk2^cmut^;ACCG* MEFs were treated with veh and Tmx respectively. C. Graph depicts percentage of cilia observed not having any breaks (i.e. normal cilia). Each dot represents an experiment with >10 observable cilia. Statistical comparison was performed by an unpaired Student’s t-test, ** denotes p<0.01. Graph depicts percentage of cilia observed being lost by the end of the experiment. Each dot represents an experiment with an initial amount of cilia >10. Statistical comparison was performed by an unpaired nonparametric Student’s t-test, * denotes p<0.05.

Larger fractions of the axoneme could also be seen breaking off the cilia (33.18%, referred to as “axonemal”) in Tmx-treated *Ttbk2^cmut^;ACCG* cells, whereas no Veh-treated cells exhibited such breakages (**Fig. 2A,B**). The net result of this is that fewer than half of the observed cilia in our live-imaged *Ttbk2^cmut^* cells + Tmx were free of axonemal breakages or budding throughout the time filmed, in contrast to more than 90% of the control cells. Moreover, we observed an average of nearly 20% of cilia lost entirely from breakages during the period of our movies in Tmx-treated *Ttbk2^cmut^* cells compared to none for Veh-treated cells (**Fig. 2C**). Therefore, our data suggest that TTBK2 maintains cilia by limiting axonemal breakages.

### TTBK2 partially maintains cilium stability by regulating actin dynamics and axonemal microtubules

Our live imaging results were reminiscent of primary cilia shedding induced by serum stimulation (16), and suggested that TTBK2 prevents breakages of the ciliary axoneme, either by promoting recruitment or retention of factors that stabilize the cilium, or by suppressing negative regulators that might promote ciliary disassembly. To address these non-mutually exclusive possibilities, we examined proteins important for stabilizing the ciliary axoneme and those that promote cilium disassembly. We examined several known regulators of cilium disassembly including phosphorylated, active AURKA, PLK1, and KIF2A (17-19) in the *Ttbk2^cmut^*cells and did not observe any accumulation of these proteins at the basal body or centrosome following loss of TTBK2 (**Fig. S2**), suggesting that TTBK2 regulates ciliary stability primarily through other mechanisms.

A major player in ciliary axonemal stability are microtubules. Specifically, the microtubules that comprise the ciliary axoneme are highly modified, particularly by acetylation and polyglutamylation, and these modifications ensure axonemal stability (20). Because we saw fragments of the axoneme of variable lengths breaking away from cilia, we assessed whether loss of these post-translational modifications could explain how these cilia are destabilized by loss of TTBK2. We found that while tubulin acetylation in the remaining cilia of the *Ttbk2^cmut^* cells was unaltered at 72 hours post-Tmx (**Fig. 3A,B**), polyglutamylation was substantially reduced relative to control cells, both in terms of the extent of polyglutamylated tubulin along the axoneme and staining intensity (**Fig. 3A,C,D**). This is consistent with what we previously observed in *Ttbk2* hypomorphic mutants (11), further suggesting that TTBK2 plays a role in ensuring that the axonemal microtubules are polyglutamylated and that this polyglutamylation may limit cilia fragmentation.

**Figure 3:**
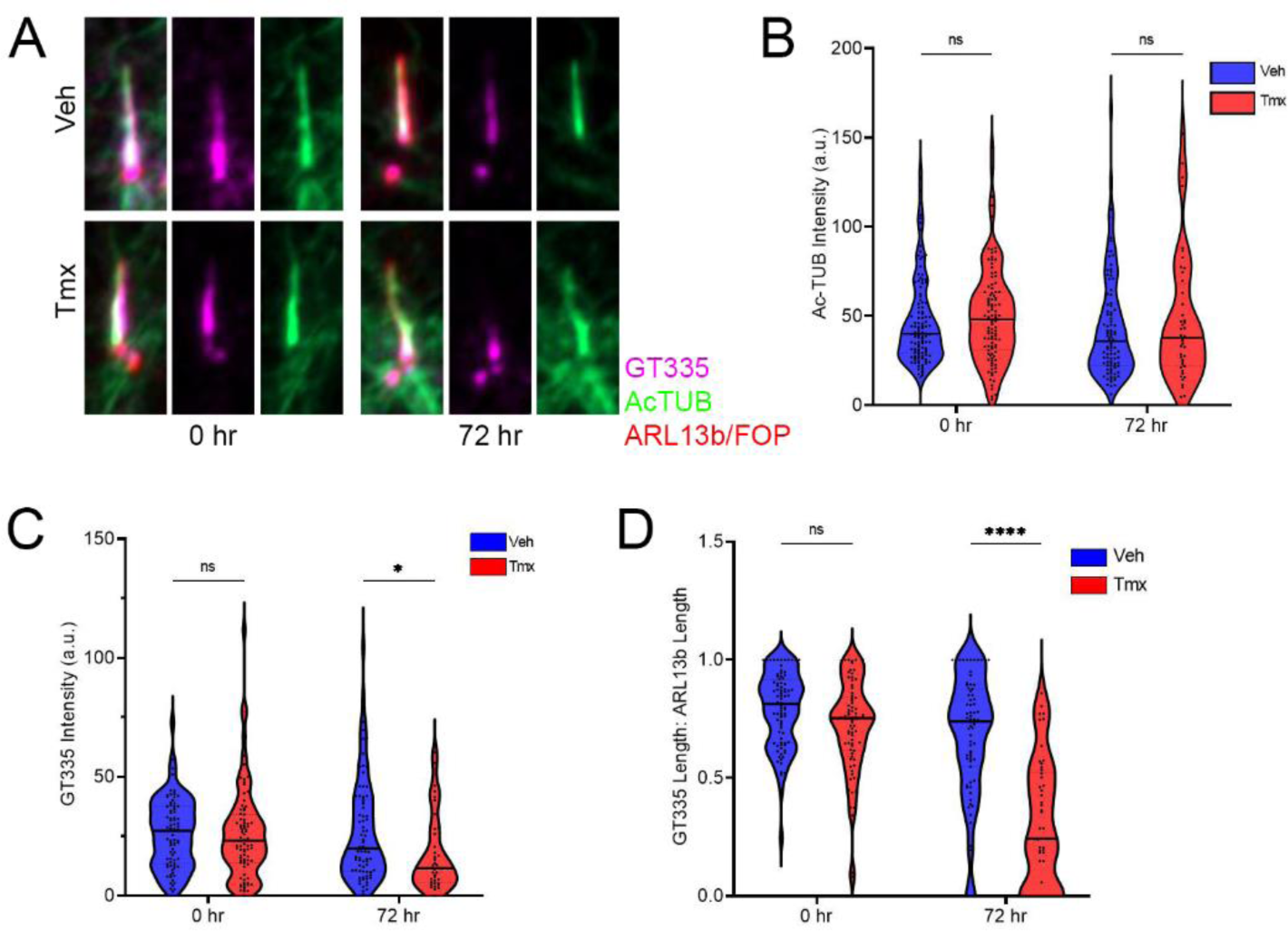
Axonemal polyglutamylation is reduced upon *Ttbk2* deletion. A. Representative images of *Ttbk2^cmut^* MEFs treated with Veh or Tmx for 0 and 72 hours and stained for AcTUB (green), GT335 (magenta) to label polyglutamylated tubulin, and ARL13B and FOP (red) to label cilia and centrosomes, respectively. B. Graph depicts violin plot of the AcTUB intensity. Each dot represents a single measurement. Results are pooled from three independent experiments. Veh 0 hr, n=114; Veh 72 hr, n=94; Tmx 0 hr, n=104; Tmx 72 hr, n=44. Statistical comparison was performed by two-way ANOVA with Tukey’s multiple comparisons test, ns denotes not significant,. C. Graph depicts violin plot of GT335 intensity along the axoneme marked by ARL13B. Each dot represents a single measurement. Results are pooled from three independent experiments. Veh 0 hr, n=79; Veh 72 hr, n=76; Tmx 0 hr, n=83; Tmx 72 hr, n=38. Statistical comparison was performed by two-way ANOVA with Tukey’s multiple comparisons test, ns denotes not significant, * denotes p<0.05. D. Graph depicts violin plot of the ratio of GT335 length to ARL13b length. Each dot represents a single measurement. Results are pooled from three independent experiments. Veh 0 hr, n=81; Veh 72 hr, n=75; Tmx 0 hr, n=76; Tmx 72 hr, n=50. Statistical comparison was performed by two-way ANOVA with Tukey’s multiple comparisons test, ns denotes not significant, **** denotes p<0.0001.

In addition to under-modification of tubulin being associated with instability of the ciliary axoneme (21), alterations to actin dynamics are also linked to ciliary breakage and shedding (15,22). We therefore tested several small molecules inhibitors that modulate specific regulators of cytoskeletal dynamics for their ability to affect the phenotype of *Ttbk2^cmut^* cells treated with Tmx, either with respect to the timing or the severity of cilia loss. These drugs included the inhibitors 3,5-bis(trifluoromethyl)pyrazole (BTP2, a drebrin inhibitor) (23), Blebbistatin (a Myosin II inhibitor) (24), Tubastatin A (TBSA, an HDAC6 inhibitor), 2,4,6 Triiodophenol (TIP, a Myosin VI inhibitor), and Cytochalasin D (CytoD, an actin polymerization inhibitor). In these experiments, we used a similar scheme to that used for our evaluation of cilia loss in *Ttbk2^cmut^* cells: cells were cultured in low-serum media for 15 hours to induce cilium assembly, followed by treatment of the cells with Tmx or Veh to induced Cre-mediated recombination, and with the indicated drugs or DMSO vehicle (**Fig. 4A**), with the start of Tmx/drug treatment defined as T0. We assessed the proportion of ciliated cells at 48 hours, since this is the time point when cilia are appreciably diminished in frequency in *Ttbk2^cmut^* cells + Tmx but are not absent as at 72 hours (**Fig 1B,C**). In control cells, treatment with the drugs had no impact on the frequency of cilia, with the exception of the Myosin VI inhibitor TIP, which modestly reduced cilia frequency (**Fig. 4B,C**). We also observed an increase in ciliary length in control cells treated with CytoD and TBSA, consistent with the known roles of actin polymerization and HDAC6 in limiting cilia length and promoting disassembly (25-28)(**Fig. 4D**).

**Figure 4:**
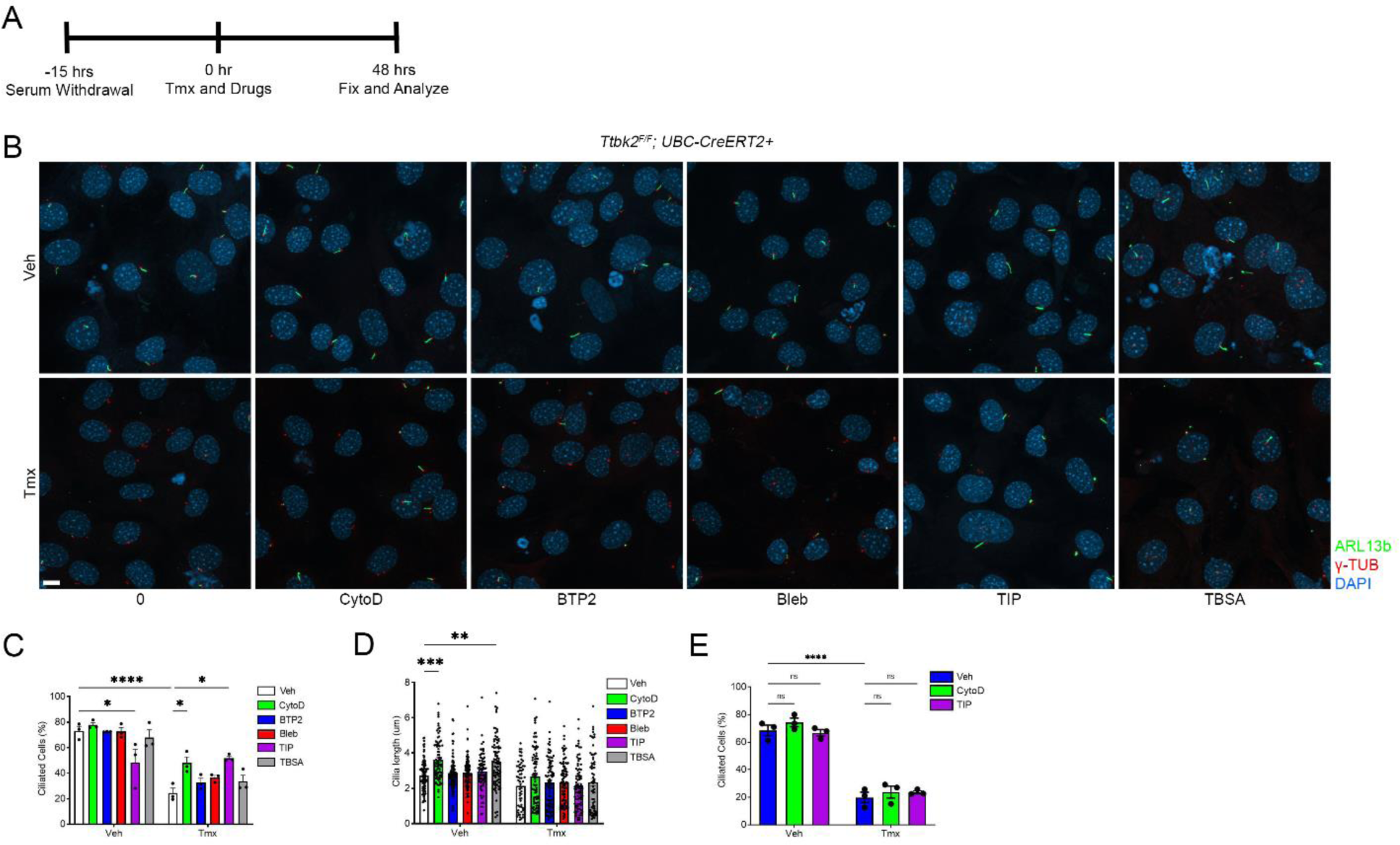
Cytochalasin D and TIP partially attenuate cilia loss due to TTBK2 deletion. A. Schematic of experimental design. *Ttbk2^cmut^* MEFs were serum starved for 15 hours before being treated with Veh or Tmx simultaneously with DMSO, CytoD (200nM), Bleb (50µ M), BTP2 (1µM), TIP (25µM), or TBSA (2µM) for 48 hours. B. Representative images of *Ttbk2^cmut^* MEFs after being treated with Veh or Tmx with the indicated drugs for 48 hrs and stained for ARL13b (green), γ-Tubulin (red), and DAPI (blue). Scale bar: 10μm. C. Graph depicts mean percentage of ciliated cells ± SEM at 48 hours. Each dot represents an experiment with >60 cells. Statistical comparison was performed by two-way ANOVA with Tukey’s multiple comparisons test, * denotes p<0.05, **** denotes p<0.0001. D. Graph depicts mean length of cilia ± SEM, measured at 48 hours. Results are pooled from three independent experiments. Each dot represents a cilium (n> 60). Statistical comparison was performed by two-way ANOVA with Tukey’s multiple comparisons test, ** denotes p<0.01, *** denotes p<0.001. E. Graph depicts mean percentage of ciliated cells ± SEM at 72 hours. Each dot represents an experiment with >60 cells. Statistical comparison was performed by two-way ANOVA with Tukey’s multiple comparisons test, ns denotes not significant, **** denotes p<0.0001.

In Tmx-treated *Ttbk2^cmut^* cells, co-treatment with Tmx plus CytoD or the Myosin VI inhibitor TIP resulted increased cilia frequency at 48 hours relative to treatment with DMSO, whereas the other inhibitors tested had no effect (**Fig. 4B,C**). This finding is consistent with the general role of actin polymerization in suppressing cilium assembly and promoting ciliary disassembly through budding and/or breakage (22). However, despite a higher frequency of cilia at 48 hours post Tmx treatment, by 72 hours, cilia were lost regardless of inhibition of these actin regulators (**Fig. 4E**). These data suggest that TTBK2 partially mediates primary cilia stability by interfering with actin dynamics. Loss of TTBK2 results in increased actin activity, leading to cilia fragmentation (**Fig. 2B**).

### Loss of TTBK2 results in changes to the centriolar satellites and deregulation of autophagy

Since perturbing the actin dynamics on a *Ttbk2^cmut^* background only had a transient stabilizing effect on cilia, this suggests that TTBK2 regulates other parallel processes that stabilize cilia. Inhibition of Myosin VI partially attenuated cilia loss (**Fig. 4B**). In addition to its role in the budding of ectosomes from the tips of cilia (22), Myosin VI is also linked to ciliogenesis through regulation of centriolar satellites (29). We previously undertook biotin proximity labeling (BioID) with TTBK2 to identify proteins potentially in the TTBK2 ciliogenesis pathway (15). In these studies, we found that of pericentriolar material and centriolar satellite-associated proteins, as well as actin regulators were significantly enriched among the TTBK2-proximate hits(15) (**Supplemental Table 1**). We therefore examined the localization of Myosin VI and the centriolar satellites in *Ttbk2^cmut^*MEFs treated with Tmx. Consistent with our findings that inhibition of Myosin VI can delay the loss of cilia in the absence of TTBK2, we found a modest but significant accumulation of Myosin VI at the basal body in *Ttbk2^cmut^* MEFs (**Fig 5A**).

**Figure 5:**
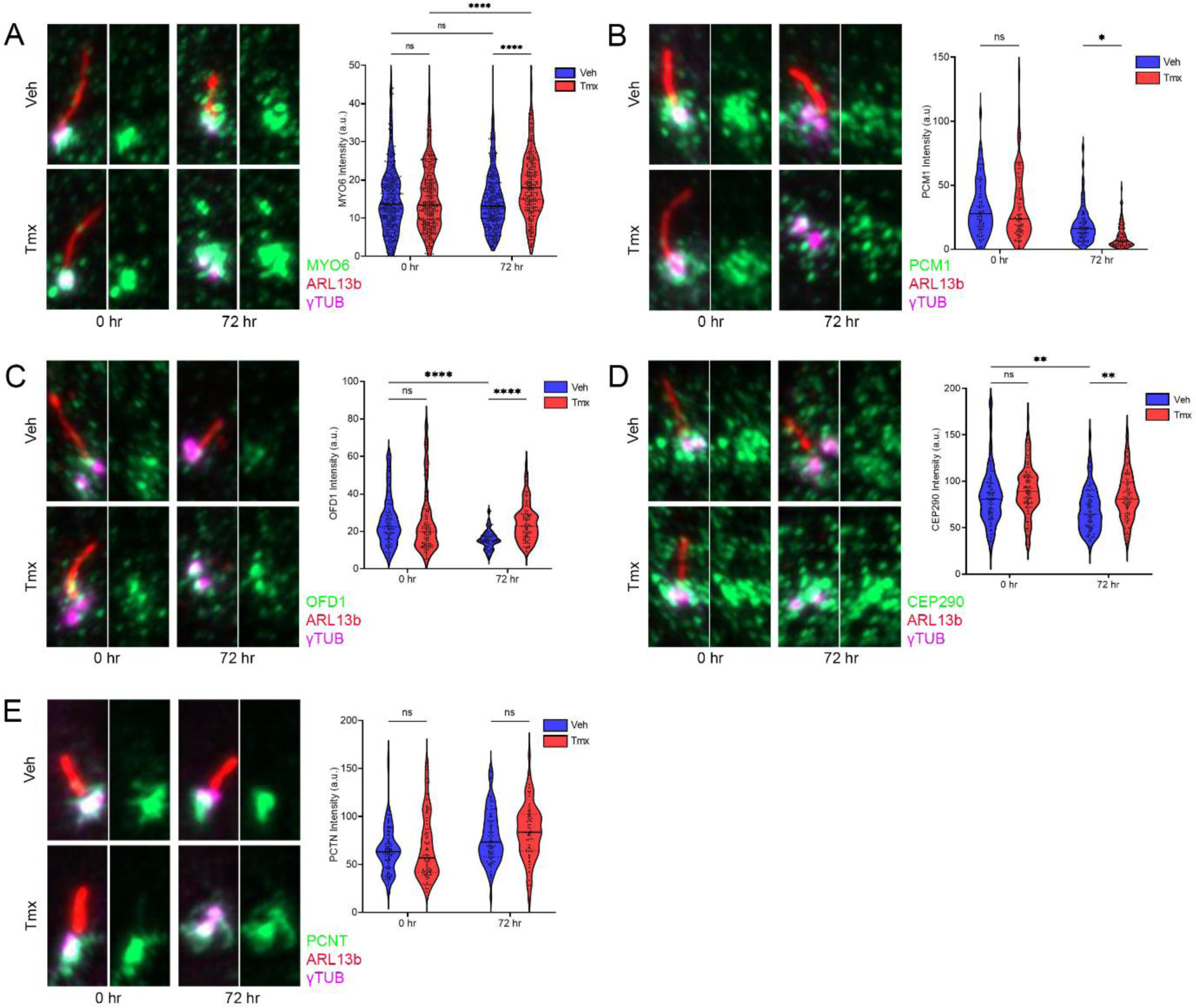
TTBK2 regulates the composition of the centriolar satellites. A. Representative images of *Ttbk2^cmut^* MEFs treated with Veh or Tmx for 0 and 72 hours and stained for MYO6 (green), ARL13b (red), and γ-Tubulin (magenta). Graph depicts mean intensity of MYO6 1μm around basal bodies. Results are pooled from three independent experiments. Each dot represents a single measurement from a basal body (n>275). Statistical comparison was performed by two-way ANOVA with Tukey’s multiple comparisons test, ns denotes not significant, * denotes p<0.05. B. Representative images of *Ttbk2^cmut^* MEFs treated with Veh or Tmx for 0 and 72 hours and stained for PCM1 (green), ARL13b (red), and γ-Tubulin (magenta). Graph depicts mean intensity of PCM1 1μm around basal bodies. Results are pooled from three independent experiments. Each dot represents a single measurement from a basal body (n>75). Statistical comparison was performed by two-way ANOVA with Tukey’s multiple comparisons test, ns denotes not significant, * denotes p<0.05. C. Representative images of *Ttbk2^cmut^* MEFs treated with Veh or Tmx for 0 and 72 hours and stained for OFD1 (green), ARL13b (red), and γ-Tubulin (magenta). Graph depicts mean intensity of OFD1 1μm around basal bodies. Results are pooled from three independent experiments. Each dot represents a single measurement from a basal body (n>75). Statistical comparison was performed by two-way ANOVA with Tukey’s multiple comparisons test, ns denotes not significant, **** denotes p<0.0001. D. Representative images of *Ttbk2^cmut^* MEFs treated with Veh or Tmx for 0 and 72 hours and stained for CEP290 (green), ARL13b (red), and γ-Tubulin (magenta). Graph depicts mean intensity of CEP290 1μm around basal bodies. Results are pooled from three independent experiments. Each dot represents a single measurement from a basal body (n>75). Statistical comparison was performed by two-way ANOVA with Tukey’s multiple comparisons test, ns denotes not significant, ** denotes p<0.01. E. Representative images of *Ttbk2^cmut^* MEFs treated with Veh or Tmx for 0 and 72 hours and stained for PCTN (green), ARL13b (red), and γ-Tubulin (magenta). Graph depicts mean intensity of PCTN at the basal bodies. Results are pooled from three independent experiments. Each dot represents a single measurement from a basal body (n>63). Statistical comparison was performed by two-way ANOVA with Tukey’s multiple comparisons test, ns denotes not significant.

Based on our findings for Myosin VI, and our BioID data linking TTBK2 to the centriolar satellites, we hypothesized that dysfunction of the satellites might contribute to the cilium stability defects we observe in *Ttbk2^cmut^* MEFs treated with Tmx. Centriolar satellites play complex roles in cilium assembly. PCM1 is required for cilia assembly (30), while other proteins such as OFD1 must be degraded at the satellites to promote cilia formation (31). We therefore assessed the composition of the centriolar satellites and the localization of key satellite proteins, including PCM1, OFD1 and CEP290 in the *Ttbk2^cmut^*cells ± Tmx. PCM1 is the main component of the centriolar satellites and serves to scaffold, assemble, and maintain the satellites (32). We found that the intensity of PCM1 was dramatically decreased by 72 hours post Tmx treatment (**Fig. 5B**), suggesting that TTBK2 might be important for the integrity of the satellites.

To determine if TTBK2 regulates other centriolar satellite components, we also assessed OFD1 and CEP290. OFD1 and CEP290, which both reside at the mother centriole in addition to the satellites, decline in intensity specifically at the satellites as serum starvation continues in control cells. However, both proteins remained elevated in *Ttbk2^cmut^*cells 72 hours following treatment with Tmx (**Fig. 5C, D**). Similar defects in centriolar satellite composition are also seen in *Ttbk2^null/null^* cells, namely a decrease in PCM1 and an increase in OFD1 and CEP290 relative to WT cells (**Fig. S3**). In contrast, PCNT, a pericentriolar material protein, was not changed by loss of TTBK2, suggesting that these defects are limited to proteins found at the centriolar satellites (**Fig. 5E**).

Mutation of a conserved aspartic acid to alanine in TTBK2’s catalytic domain blocks its kinase activity (33). While this kinase-dead TTBK2 correctly localizes to the mother centriole, it cannot rescue cilium assembly in *Ttbk2^null/null^*cells (10), demonstrating that TTBK2 kinase activity is required for its role in cilium assembly. To determine if TTBK2 simply acts as a scaffold for centriolar satellites at the cilium or if its kinase activity is required to regulate their composition, we stably expressed a kinase-dead TTBK2 or a WT TTBK2 in our *Ttbk2^cmut^*cells. Tmx-treated *Ttbk2^cmut^* cells stably expressing a kinase-dead TTBK2-GFP were unable to rescue PCM1 localization to the basal body whereas *Ttbk2^cmut^* cells stably expressing WT TTBK2-GFP had restored PCM1 levels at centriolar satellites (**Fig. S4**). Thus, TTBK2 regulates the composition of the centriolar satellites after cilium assembly through its kinase activity.

OFD1 is normally degraded at the centriolar satellites via autophagy to promote cilium assembly. In autophagy-deficient cells, OFD1 is retained at the satellites, inhibiting ciliogenesis (31). The accumulation of OFD1 at the centriolar satellites in *Ttbk2^cmut^* cells suggested a deregulation of autophagy. Mammalian target of rapamycin (mTOR) regulates a number of cellular processes including cell growth, survival, metabolism, stress responses, and autophagy (34). In nutrient-rich conditions, mTOR is phosphorylated at S2448 and inhibits autophagy (35). Compared to the start of the Tmx-induction, the ratio of phospho-mTOR to mTOR levels remained relatively unchanged in Veh-treated *Ttbk2^cmut^* cells whereas Tmx-treated *Ttbk2^cmut^*cells had a higher phospho-mTOR to mTOR ratio (**Fig. 6A**). This further supports a disruption in autophagy upon deletion TTBK2, due at least in part to mTOR activation.

**Figure 6:**
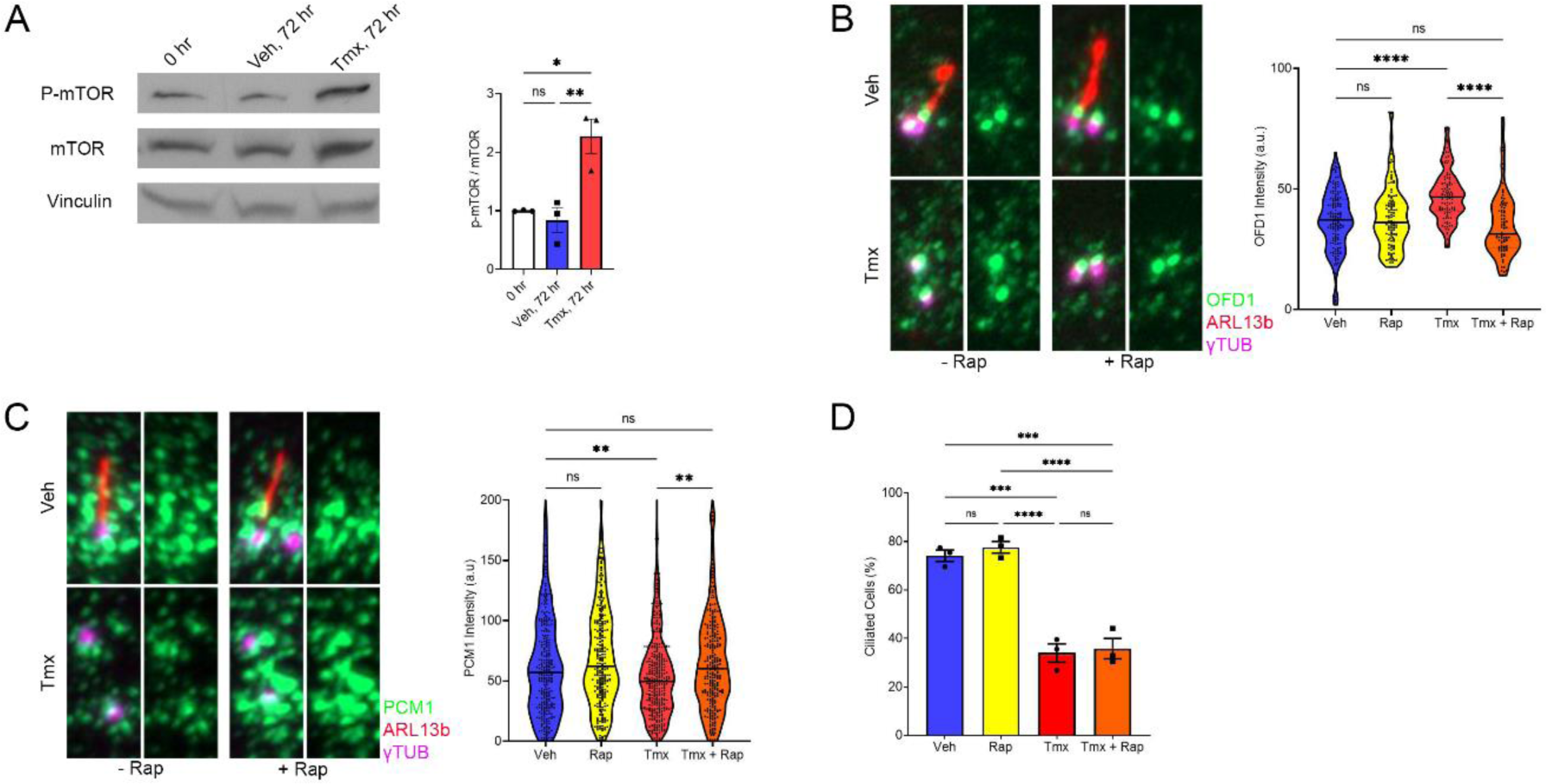
TTBK2 regulates the centriolar satellites through autophagy. A. Western blot of P-mTOR, mTOR, and Vinculin from total lysates of *Ttbk2^cmut^*MEFs pre- treatment and treated with Veh or Tmx for 72 hrs. Graph depicts densitometric analysis of p-mTOR and mTOR levels of *Ttbk2^cmut^* MEFs treated with Veh or Tmx for 72 hrs normalized to pre-treatment. Statistical comparison was performed by one-way ANOVA with Tukey’s multiple comparisons test, ns denotes not significant, * denotes p<0.05, ** denotes p<0.01. B. Representative images of *Ttbk2^cmut^* MEFs treated with Veh, Rap, Tmx, or Tmx+Rap for 72 hours and stained for OFD1 (green), ARL13b (red), and γ-Tubulin (magenta). Graph depicts mean intensity of OFD1 1μm around basal bodies. Results are pooled from three independent experiments. Each dot represents a single measurement from a basal body (n>85). Statistical comparison was performed by one-way ANOVA with Tukey’s multiple comparisons test, ns denotes not significant, **** denotes p<0.0001. C. Representative images of *Ttbk2^cmut^* MEFs treated with Veh, Rap, Tmx, or Tmx+Rap for 72 hours and stained for PCM1 (green), ARL13b (red), and γ-Tubulin (magenta). Graph depicts mean intensity of PCM1 1μm around basal bodies. Results are pooled from three independent experiments. Each dot represents a single measurement from a basal body (n>260). Statistical comparison was performed by one-way ANOVA with Tukey’s multiple comparisons test, ns denotes not significant, ** denotes p<0.01. D. Graph depicts mean percentage of ciliated cells ± SEM. Each dot represents an experiment with >50 cells. Statistical comparison was performed by one-way ANOVA with Tukey’s multiple comparisons test, ns denotes not significant, *** denotes p<0.001, **** denotes p<0.0001.

Rapamycin promotes autophagy by interfering with mTOR’s ability to bind to its effectors (34) thereby inhibiting its activity. To test whether the centriolar satellite defects we observed in *Ttbk2^cmut^*cells are due to mTOR activation leading to reduced autophagy, we treated *Ttbk2^cmut^*cells with rapamycin and assessed whether we could rescue the cilia loss or other defects caused by TTBK2 deletion. Rapamycin did not change OFD1 levels in control cells but reduced OFD1 levels in Tmx-treated cells comparable to control cells (**Fig. 6B**). Surprisingly, rapamycin also rescued PCM1 loss at the satellites in Tmx-treated cells (**Fig. 6C**). However, despite restoring PCM1 and degrading OFD1 at the satellites, rapamycin was unable to rescue cilia in Tmx-treated *Ttbk2^cmut^* cells (**Fig. 6D**). Our data suggest that while TTBK2 regulates the centriolar satellite composition through autophagy, these defects do not fully drive the loss of cilia that occurs in the absence of TTBK2.

### TTBK2 is required for pools of IFT proteins at the basal body that are essential to maintain the ciliary axoneme

Thus far, our data suggest that loss of TTBK2 destabilizes the cilium through the effects of both perturbed actin dynamics and dysregulation of the centriolar satellites. While the centriolar satellite composition is disrupted due to *Ttbk2* deletion, restoration of PCM1 and OFD1 to levels comparable to control cells with rapamycin was not sufficient to rescue the cilia loss phenotype (**Fig. 6D**). The centrosomal compartment itself, where PCTN and the cilia disassembly proteins localize, remains unchanged in the Tmx-treated *Ttbk2^cmut^* cells (**Fig. 5E** and **Fig. S2**). However, the axoneme exhibited dramatic changes such as increased cilia fragmentation (**Fig. 2A-C**) and reduced axonemal polyglutamylation (**Fig. 3A-C**). This suggests that loss of axonemal integrity may drive cilia loss in Tmx-treated *Ttbk2^cmut^* cells, and we further reasoned that this loss of axonemal integrity may be due to disrupted ciliary trafficking. Ciliary trafficking is governed by IFT proteins (9,36). Our previous work demonstrated that TTBK2 is required for the recruitment of IFT proteins to the mother centrioles prior to the assembly of the ciliary axoneme (10).

We therefore examined the localization of the IFTB component IFT88 and the IFTA component IFT140 to both the basal body pool and the ciliary axoneme in *Ttbk2^cmut^* cells ± Tmx over the 72-hour time course. We found that for both IFT88 and IFT140, the percentage of cells with IFT protein at the basal body declines within 48 hours of Tmx treatment (**Fig. 7A, C**). This also corresponds with when cilia begin to disappear in the *Ttbk2^cmut^* cells treated with Tmx (**Fig. 1C**).

**Figure 7:**
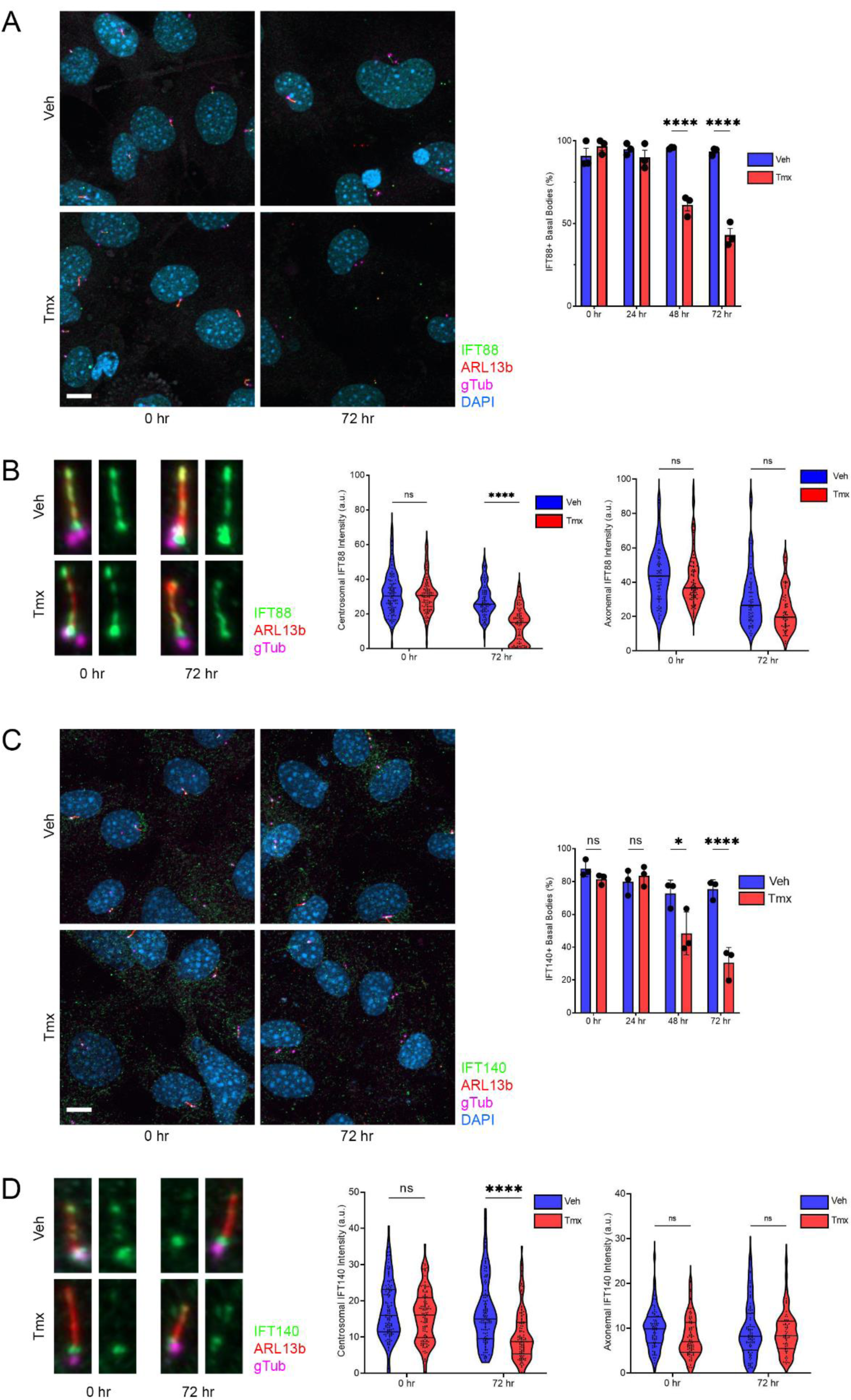
TTBK2 maintains the basal body pools of IFT. A. Representative images of *Ttbk2^cmut^* MEFs treated with Veh or Tmx for 0 and 72 hours and stained for IFT88 (green), ARL13b (red), γ-Tubulin (magenta), and DAPI (blue). Graph depicts mean percentage of basal bodies with IFT88 ± SEM. Each dot represents an experiment with >50 basal bodies. Scale: 10μm. Statistical comparison was performed by two-way ANOVA with Tukey’s multiple comparisons test, **** denotes p<0.0001. B. Representative images of *Ttbk2^cmut^* MEFs treated with Veh or Tmx for 0 and 72 hours and stained for IFT88 (green), ARL13b (red), and γ-Tubulin (magenta). Graph depicts mean intensity of IFT88 at basal bodies. Results are pooled from three independent experiments. Each dot represents a single measurement from a basal body (n>75). Statistical comparison was performed by two-way ANOVA with Tukey’s multiple comparisons test, ns denotes not significant, **** denotes p<0.0001. Graph depicts mean intensity of IFT88 along the axoneme. Results are pooled from three independent experiments. Each dot represents a single measurement from a basal body. Veh 0 hr, n=76; Veh 72 hr, n=72; Tmx 0 hr, n=95; Tmx 72 hr, n=57. Statistical comparison was performed by two-way ANOVA with Tukey’s multiple comparisons test, ns denotes not significant. C. Representative images of *Ttbk2^cmut^* MEFs treated with Veh or Tmx for 0 and 72 hours and stained for IFT140 (green), ARL13b (red), γ-Tubulin (magenta), and DAPI (blue). Graph depicts mean percentage of basal bodies with IFT88 ± SEM. Each dot represents an experiment with >50 basal bodies. Scale: 10μm. Statistical comparison was performed by two-way ANOVA with Tukey’s multiple comparisons test, * denotes p<0.05, **** denotes p<0.0001. D. Representative images of *Ttbk2^cmut^* MEFs treated with Veh or Tmx for 0 and 72 hours and stained for IFT140 (green), ARL13b (red), and γ-Tubulin (magenta). Graph depicts mean intensity of IFT140 at basal bodies. Results are pooled from three independent experiments. Each dot represents a single measurement from a basal body (n>75). Statistical comparison was performed by two-way ANOVA with Tukey’s multiple comparisons test, ns denotes not significant, **** denotes p<0.0001. Graph depicts mean intensity of IFT140 along the axoneme. Results are pooled from three independent experiments. Each dot represents a single measurement from a basal body. Veh 0 hr, n=77; Veh 84 hr, n=72; Tmx 0 hr, n=89; Tmx 72 hr, n=61. Statistical comparison was performed by two-way ANOVA with Tukey’s multiple comparisons test, ns denotes not significant

Analysis of fluorescence intensity revealed that at 72 hours post Tmx treatment, the *Ttbk2^cmut^* cells showed diminished levels of both IFT88 and IFT140 in the basal body pools. By contrast, the intensity of both IFT proteins within the remaining ciliary axonemes was not significantly affected (**Fig. 7B, D**). This suggests that the basal body pool of IFT proteins that plays a critical role in maintaining the integrity of the ciliary axoneme, and that TTBK2 is required to maintain this pool following cilium assembly, in addition to being required to recruit IFT proteins during ciliogenesis.

Our findings suggest that TTBK2 regulates primary cilia stability through several parallel pathways (**Fig. 8**). TTBK2 prevents cilia fragmentation by controlling actin dynamics after ciliogenesis. It also regulates the composition of the centriolar satellites through its kinase activity and autophagy. Finally, in addition to being required for IFT recruitment during ciliogenesis, TTBK2 is required for the retention or replenishment of IFT pools at the basal body. These findings reveal multiple mechanisms by which the primary cilium is maintained after being assembled.

**Figure 8:**
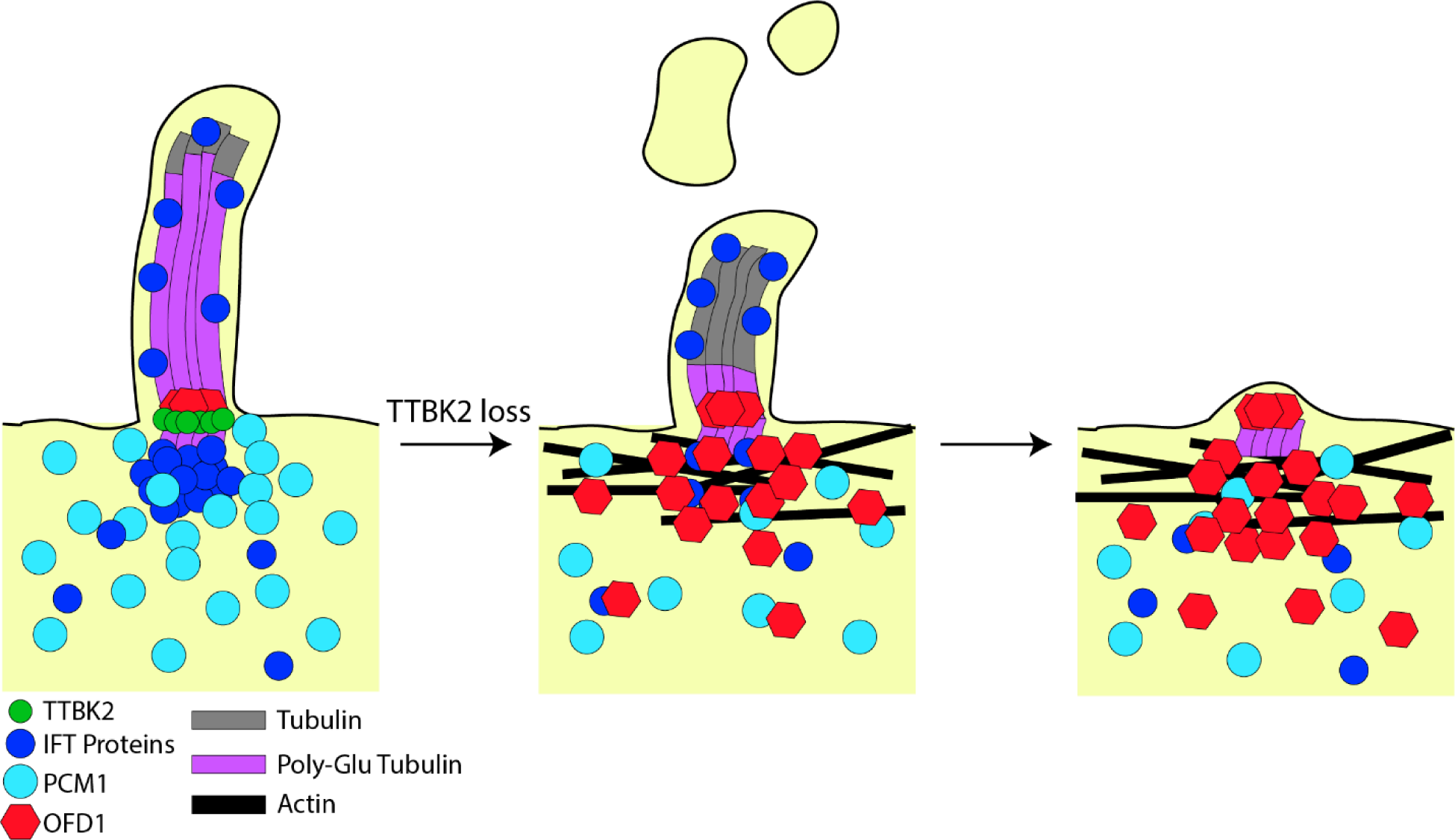
TTBK2 maintains primary cilia through multiple pathways. Depletion of TTBK2 leads to a progressive loss of IFT proteins and PCM1 at the basal body and an accumulation of OFD1 and actin filaments. This leads to reduced polyglutamylation of the axoneme and increased fragmentation of primary cilia resulting in eventual primary cilia loss.

## Discussion

In this study, we use an inducible genetic system to assess the requirements for TTBK2 in maintaining cilia structure. Using these tools, we were able to perturb cilia after cells had undergone ciliogenesis through tamoxifen-induced deletion of TTBK2. This has allowed us to assess the dynamic changes to primary cilia as TTBK2 is lost and uncover additional requirements for TTBK2 in regulating primary cilia.

Our previous work on TTBK2 suggested that TTBK2 regulates primary cilia structure and stability, but the precise mechanisms by which TTBK2 does this were left unknown. Serum re-addition following the induction of cilia by serum-withdrawal results in cilia loss in cultured fibroblasts that correlates with TTBK2 loss or removal from the basal body (10). Cells with hypomorphic mutations in TTBK2 form unstable cilia (11). Finally, deletion of TTBK2 in adult mice results in reduced cilia frequency in the adult brain as rapidly as 15 days following Tmx induction in *Ttbk2^cmut^* mice. In this study, we visualize how cilia loss occurs in the absence of TTBK2 through live imaging. While a low frequency of ciliary excision from the tip is proposed to be a mechanism for maintaining ciliary length (37), both the proportion of cilia that form buds and the frequency of budding events per cilium increases when TTBK2 is depleted. Additionally, some cilia shed larger fragments of the axoneme upon TTBK2 loss, reminiscent of events linked to cilia disassembly, where cilia are lost predominantly through whole cilium shedding (16).

Ciliary excision and whole cilium shedding can be attenuated using actin inhibitors and HDAC6 inhibition respectively (16,22). The similarities between these studies and our observations prompted us to assess whether TTBK2 maintains primary cilia by regulating actin dynamics and/or suppressing the cilia disassembly pathway. *Ttbk2* deletion does not alter the localization of the cilia disassembly proteins KIF2A, PLK1, or phospho-AURKA at the basal body, suggesting that TTBK2 does not principally regulate cilia maintenance by suppressing cilia disassembly proteins. Corroborating this finding, we also show that treatment of *Ttbk2^cmut^* cells with the HDAC6 inhibitor TBSA did not lead to rescue in cilia frequency at any point in our time-course.

We treated Tmx-treated *Ttbk2^cmut^* cells with other small molecule inhibitors previously found to suppress primary cilia excision. Only CytoD and TIP partially rescued cilia loss. CytoD inhibits actin polymerization and TIP inhibits myosin VI, a motor protein that directs movement towards the minus end of actin filaments, suggesting that TTBK2 may mediate primary cilia stability by regulating actin dynamics. In addition to its role in regulating cilium assembly, TTBK2 is also linked to cytoplasmic microtubule dynamics (38,39). and recent studies point to crosstalk between microtubules and actin filaments (40,41). Recent work from our lab identified CSNK2A as a direct interactor of TTBK2 that negatively modulates TTBK2 function at the ciliary base(15). In CSNK2A knockout cells, cilia are unstable, undergoing frequent excision events, and actin and actin modulators are enriched at the ciliary tip, associated with those excision events (15). Our TTBK2 BioID screen also identified several actin regulators as TTBK2-proximate proteins (15). These data suggest that TTBK2 mediates cilia stability in part by inhibiting actin polymerization. *Ttbk2* deletion relieves this inhibition and cytochalasin D restores it, albeit temporarily attenuating cilia loss, due to additional requirements for TTBK2 in cilia initiation and maintenance.

Consistent with TIP temporarily attenuating cilia loss resulting from TTBK2 deletion, Myosin VI levels increased at the centrioles in Tmx-treated *Ttbk2^cmut^* cells. In addition to its role in trafficking proteins along actin filaments, Myosin VI has also been reported to regulate the centriolar satellite composition (29). Recent evidence also suggests that the centriolar satellite scaffold PCM1 is also involved in cilia maintenance (42). The centriolar satellites were accordingly disrupted as a function of *Ttbk2* deletion with PCM being lost and OFD1 and CEP290 being retained or accumulating. TTBK2’s kinase activity is required for ciliogenesis and the maintenance of PCM1 at the basal body, but whether TTBK2 directly phosphorylates PCM1 is not currently known.

In addition to PCM1, TTBK2 also regulates the levels of OFD1 and CEP290 at the satellites and basal body. While TTBK2 maintains PCM1 levels at the satellites, it appears to regulate the degradation of OFD1 and CEP290 at the satellites. Prior work has shown that while OFD1 recruitment to the centriole is required for the maturation of the mother centriole and therefore the initiation of ciliogenesis (7), but its degradation by autophagy at the satellites is also required for cilium assembly (31). Serum starvation induces autophagy (43), and autophagy promotes the degradation of OFD1 at the satellites (31). Our data suggest that cells are continuing to undergo autophagy during serum withdrawal even after ciliogenesis is underway. Our data reveal that TTBK2 is involved in this process, since OFD1 accumulates when TTBK2 is lost after cilium assembly. Rapamycin-induced autophagy restores OFD1 to levels comparable to those of control cells, suggesting that the mTOR-mediated autophagy is linked to OFD1 degradation, and that TTBK2 inhibits mTOR activation. The precise mechanism by which TTBK2 regulates OFD1 degradation will need to be investigated in the future.

PCM1 and OFD1 colocalize to centriolar satellites, and their interaction is well established (31,42,44). PCM1 is proposed to mediate OFD1 degradation by facilitating its interaction with LC3 which targets OFD1 for degradation (31). Interestingly, rapamycin-induced autophagy is also able to restore PCM1 levels around the basal body suggesting that autophagy not only promotes the degradation of OFD1 but also maintains PCM1 levels at the satellites-perhaps even regulating the composition of the centriolar satellites as a whole. Our BioID screen also identified several autophagy-related proteins as proximate to TTBK2, suggesting that TTBK2 may interact with these proteins. However, the link between TTBK2 and regulation of autophagy remains to be explored.

Neither restoration of the centriolar satellites by inducing autophagy, nor disrupting actin polymerization fully rescued the cilia loss that results from *Ttbk2* deletion, leading us to assess IFT. IFT is required not only for cilium assembly but also to maintain cilia structure through continuous trafficking. Tamoxifen-induced deletion of *Ift88* in the kidneys and in the brain leads to a loss of primary cilia in adult mice (12,45). Recent work suggests that IFT is recruited to the basal body by a diffusion-to-capture mechanism (46), though how the basal body “captures” IFT remains unknown. Our work reveals that in addition to its role in IFT recruitment during cilium assembly, TTBK2 is also indispensable for IFT maintenance after cilium assembly. Our results suggest that TTBK2 is involved in modifying the basal body such that it continually replenishes IFT. In the absence of TTBK2, the basal body IFT pools therefore become depleted and ultimately disappear. The mother centriole’s distal appendages are required for TTBK2 recruitment (6,8). In addition, several recent studies have shown that TTBK2 phosphorylates some of these distal appendage proteins including CEP83, CEP89 and CEP164 (47,48) perhaps modifying it in a way that allows for the capture of IFT proteins.

In sum, our study supports a model in which TTBK2 acts via to multiple mechanisms to regulate cilium initiation and maintenance, underscoring the central role of this kinase in regulating primary cilia. We propose that TTBK2 regulates the centrosomal compartment of primary cilia which contributes to building and maintaining stable primary cilia. Our data suggest that TTBK2 regulates the actin network around the basal body and in parallel regulates the centriolar satellites via autophagy. *Ttbk2* deletion causes dysregulation of the centrosomal compartment leading to loss of axonemal integrity and its fragmentation. In a separate pathway, TTBK2 is also regulating IFT recruitment that maintains the axonemal composition and length, lost when TTBK2 is depleted.

## Acknowledgements

We thank Duke University Proteomics and Metabolomics Core facility. This work was supported by NIH R01 HD099784 and R03 TR003392 to SCG, and by and NSF graduate research fellowship to AN. We thank Don Fox and members of the Goetz lab for critical comments on this work.

## Materials and Methods

### Ethics statement

The use and care of mice as described in this study was approved by the Institutional Animal Care and Use Committee Duke University (approval numbers A218-17-09 and A175-20-08). Euthanasia for the purpose of harvesting embryos was performed by cervical dislocation, and all animal studies were performed in compliance with internationally accepted standards.

### Mouse Strains

Mouse strains used in this study were *Ttbk2^loxP/loxP^*(*Ttbk2^f/f^*) mice generated in a previous study (12), *UBC-CreERT2* mice acquired from Jackson Laboratory (#007001), and *Arl13b-* mCherry*;Centrin2-GFP* mice kindly donated from KV Anderson (14).

### Cell culture

*Ttbk2F/F; UBC-CreERT2+* mouse embryonic fibroblasts (MEFs) were derived from embryonic day (E) 10.5 mouse embryos and cultured in Dulbecco’s Modified Eagle Medium supplemented with 10% fetal bovine serum and 1% penicillin/streptomycin (cDMEM) at 5% CO_2_ at 37°C. Unless indicated otherwise, cells were plated on Gelatin-coated coverslips and serum starved for 15 hours in cDMEM. Cells were then treated with 1µM 4-hydroxytamoxifen (Tmx) for the indicated duration.

### Transient and Stable Transfection

MEFs were transfected using Lipofectamine 2000 (ThermoFisher 11668) in OptiMEM (Gibco 31985). For stable expression of TTBK2-GFP, retroviruses were produced and used to infect MEFs according to standard protocols. MEFs were then put under 500µg/mL G418 selection. Plasmids used were pFLAP-Dest-TTBK2 described previously (10).

### Drug and inhibitor treatment

Drugs were added to cells at the time of Tmx induction at the following concentrations: Cytochalasin D (Santa Cruz SC-201442) (200nM in DMSO), Blebbistatin (VWR S7099) (50µM in DMSO), BTP2 (Abcam ab144413) (1µM in DMSO), Rapamycin (Cell Signaling 99045) (25nM in DMSO), Tubastatin A (Tocris 6270) (2µM), and TIP (Sigma 137723) (25µM).

### Immunofluorescence

Cells were then fixed in 4% paraformaldehyde for 5 minutes and then ice-cold methanol for 5 minutes, washed 3x with PBS for 5 minutes each, and blocked with PBS + 5% goat serum + 1% BSA + 0.1% Triton X-100 (blocking buffer) for 30 minutes. Next, cells were incubated with primary antibodies diluted in blocking buffer overnight at 4°C, washed 3x with PBS for 5 minutes each, incubated with secondary antibodies diluted in blocking buffer for 30 minutes at room temperature. Finally, cells were stained with DAPI in PBS for 5 minutes, washed two additional times with PBS for 5 minutes each, and mounted onto glass slides using Prolong Gold (Invitrogen P36930).

### Antibodies

Primary antibodies used were mouse anti-ARL13b (NeuroMab 75-287), rabbit anti-ARL13b (Proteintech 17711-1-AP), rabbit anti-phospho-Aurora A (Cell Signaling 3079), rabbit anti-CEP290 (Proteintech 22490-1-AP), rabbit anti-GFP (Invitrogen A11122), rabbit anti-FOP (Proteintech 11343-1-AP), mouse anti-GFP (Proteintech 66002-1-Ig), rabbit anti-IFT88 (Proteintech-11744), rabbit anti-IFT140 (Proteintech 17460-1-AP), rabbit anti-KIF2A (Abcam ab37005), rabbit anti-OFD1 (kindly donated from JF Reiter), rabbit anti-mTOR (Cell Signaling 2971), rabbit anti-phospho-mTOR (Cell signaling 2983), rabbit anti-PCM1 (Proteintech 19856-1-AP), rabbit anti-PCNT (Biolegend 923701), mouse anti-PLK1 (Thermofisher 33-1700), rabbit anti-TTBK2 (Sigma HPA018113), mouse anti-γ-Tubulin (Sigma-Aldrich T6557).

### Microscopy

Immunofluorescent images were taken using a Zeiss AxioObserver wide field microscope equipped with an Axiocam 506mono camera and Apotome.2 optical sectioning with structured illumination. Z-stacks were taken at 0.5μm intervals and maximum intensity projections were then created. Image processing and quantifications were performed using ImageJ. In brief, cilia frequency was quantified by counting the number of ARL13b+ cells vs DAPI+ cells. For intensity measurements, background was subtracted, and the mean intensity was recorded for a drawn region of interest. For centriolar satellite proteins, a 1μm ROI was made around the basal body and the intensity was then measured.

### Western Blotting

*Ttbk2^cmut^* MEFs were plated in a 10cm plate and allowed to reach confluency before being serum starved for 15 hrs. Cells were then given Veh or Tmx for 0hr or 72hr and lysed in RIPA buffer (150mM NaCl, 1% NP-40, 0.5% Sodium Deoxycholate, 0.1% SDS, 50mM Tris pH 8.0). Protein concentration was measured using a BCA assay and samples were boiled at 95C for 5min to denature proteins. Total lysates were separated by SDS-PAGE and transferred onto a PVDF membrane. The membrane was blocked with 5% milk in TBST for 1hr at RT, incubated with primary antibodies in blocking buffer overnight at 4C, and incubated with HRP-conjugated secondary antibodies for 1hr at RT. Membranes were developed with an ECL. Western blots were analyzed using densitometric analysis in FIJI.

### Time-lapse Imaging

*Ttbk2^cmut^;ACCG* MEFs were seeded in an 8-well μ-Slide (IBIDI 80826), serum starved, and treated with ethanol or TMX. The cells were then imaged at 15min intervals from 30 hrs to 50 hrs post-Tmx induction in an incubation chamber maintained with 5% CO_2_ at 37°C by a Pecon CO_2_ Module S and TempModule S. The recorded cilia were then analyzed for abnormalities and loss using Zen 2.0.

### Quantification and Statistics

Data are reported as arithmetic means ± SEM. Statistical analysis was done with GraphPad Prism 8. Most experiments were analyzed using a 2-way ANOVA and a Tukey post hoc test.

## Legends to Figures

**Figure S1:**
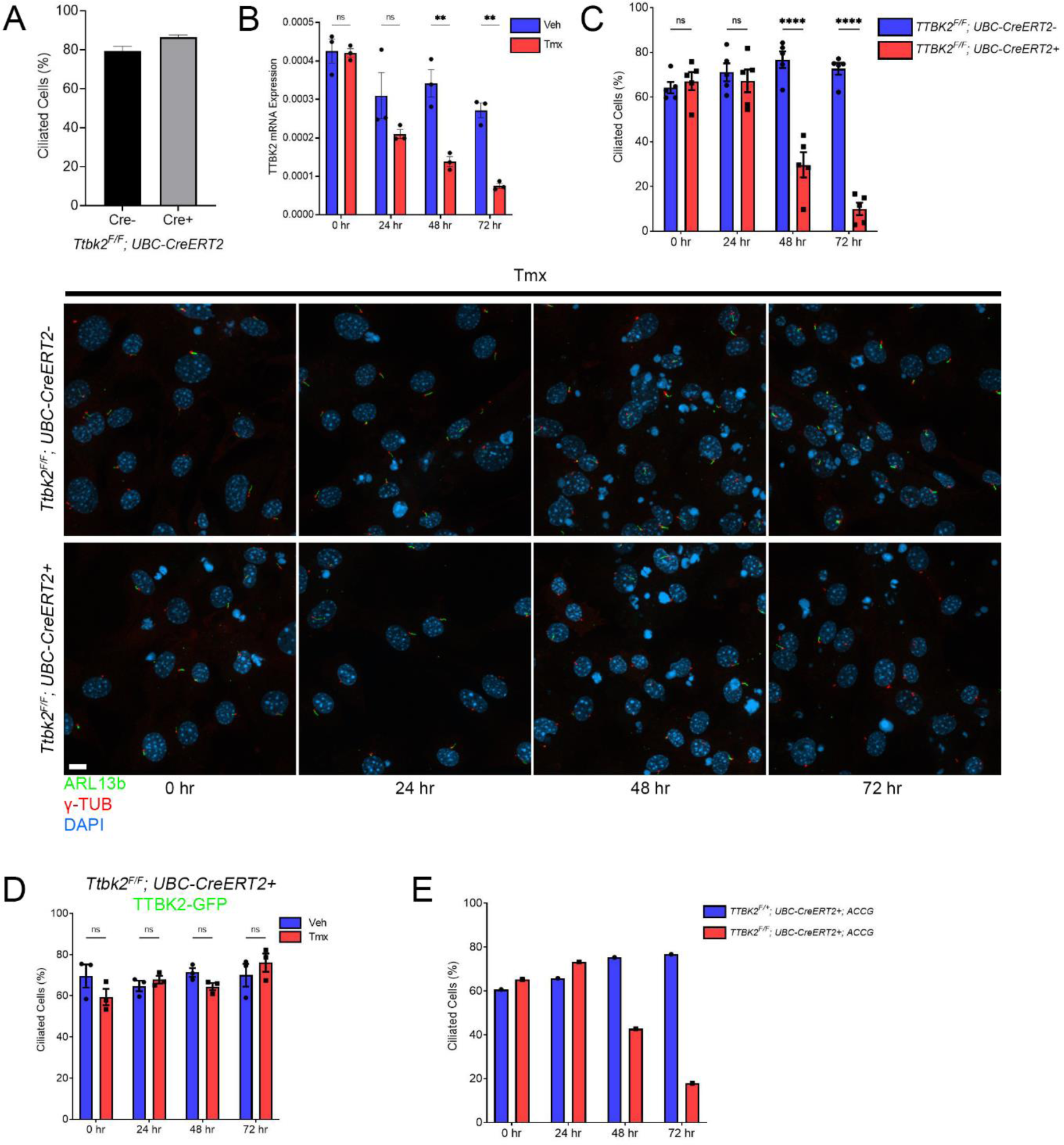
Tmx induces Cre-mediated deletion of *Ttbk2fl/fl*. A. Mean percentage of ciliated cells in *Ttbk2fl/fl; UBC-CreERT2+* and *Tbk2fl/fl; UBC-CreERT2-*serum starved for 48 hours. B. Graph depicts TTBK2 mRNA expression levels in *Ttbk2fl/fl; UBC-CreERT2+* treated with Veh or Tmx at 0, 24, 48, and 72 hours. Statistical comparison was performed by two-way ANOVA with Tukey’s multiple comparisons test, ns denotes not significant, ** denotes p<0.01. C. Representative images of *Ttbk2fl/fl; UBC-CreERT2-* and *Ttbk2fl/fl; UBC-CreERT2+* MEFs treated with Tmx for 0, 24, 48, and 72 hours and stained for ARL13b (green), γ-Tubulin (red), and DAPI (blue). Scale bar: 10μm. Graph depicts mean percentage of ciliated cells ± SEM. Each dot represents an experiment with >50 cells. Statistical comparison was performed by two-way ANOVA with Tukey’s multiple comparisons test, ns denotes not significant, **** denotes p<0.0001. D. *Ttbk2fl/fl; UBC-CreERT2+* MEFs expressing TTBK2-GFP were treated with Tmx. Graph depicts mean percentage of ciliated cells ± SEM. Each dot represents an experiment with >50 cells. Statistical comparison was performed by two-way ANOVA with Tukey’s multiple comparisons test, ns denotes not significant,. E. *Ttbk2fl/+; UBC-CreERT2+;ACCG* and *Ttbk2fl/fl; UBC-CreERT2+;ACCG* MEFs expressing TTBK2-GFP were treated with Tmx. Graph depicts percentage of ciliated cells ± SEM. Each dot represents an experiment with >70 cells.

**Figure S2:**
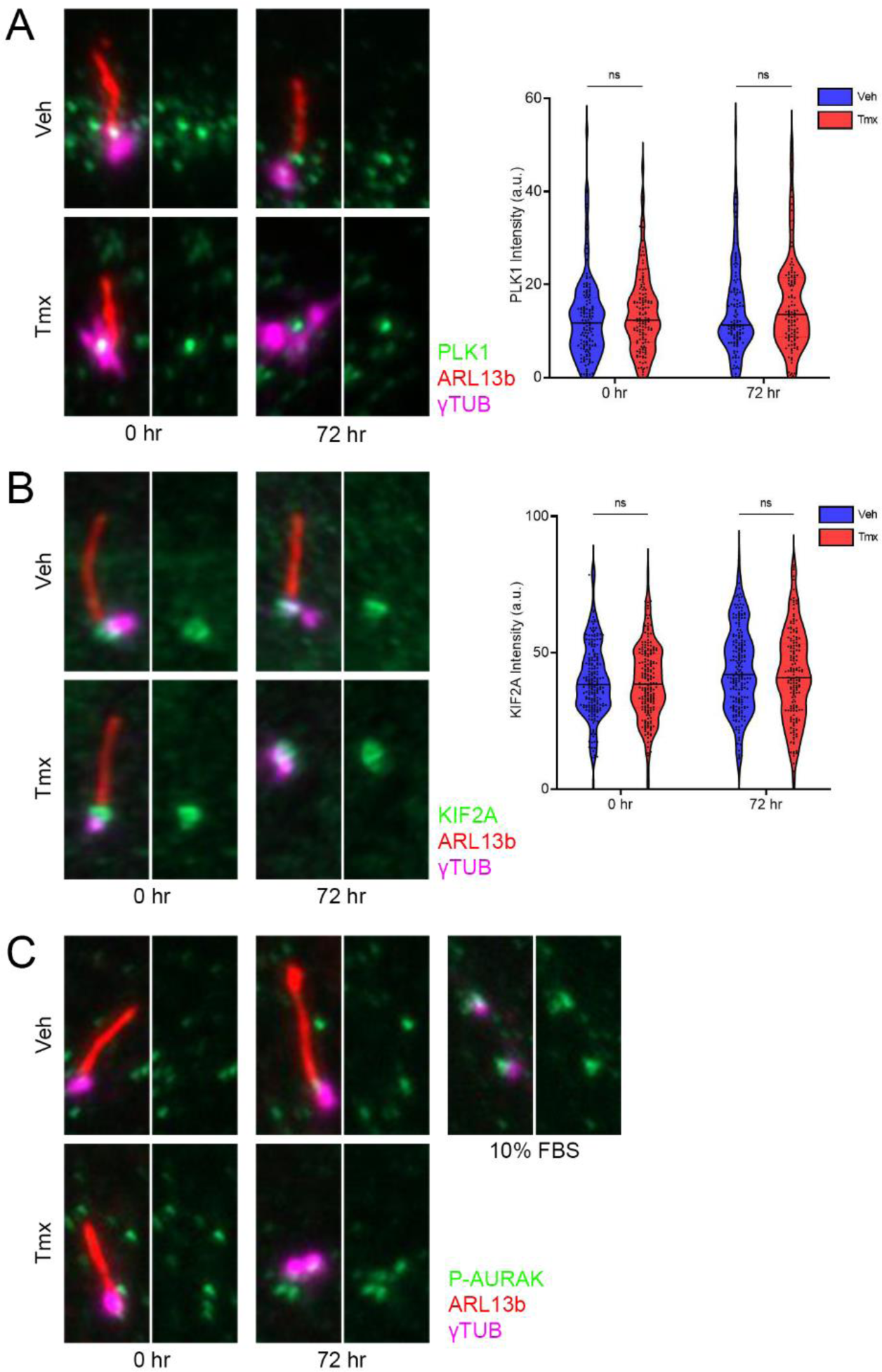
TTBK2 does maintain cilia by limiting the recruitment of cilia disassembly factors. A. Representative images of *Ttbk2^cmut^* MEFs treated with Veh or Tmx for 0 and 72 hours and stained for PLK1 (green), ARL13b (red), and γ-Tubulin (magenta). Graph depicts mean intensity of PLK1 at the basal bodies. Results are pooled from three independent experiments. Each dot represents a single measurement from a basal body (n>112). Statistical comparison was performed by two-way ANOVA with Tukey’s multiple comparisons test, ns denotes not significant. B. Representative images of *Ttbk2^cmut^* MEFs treated with Veh or Tmx for 0 and 72 hours and stained for KIF2A (green), ARL13b (red), and γ-Tubulin (magenta). Graph depicts mean intensity of KIF2A at the basal bodies. Results are pooled from three independent experiments. Each dot represents a single measurement from a basal body (n>164). Statistical comparison was performed by two-way ANOVA with Tukey’s multiple comparisons test, ns denotes not significant. C. Representative images of *Ttbk2^cmut^* MEFs treated with Veh or Tmx for 0 and 72 hours and *Ttbk2^cmut^* MEFs cultured in 10% FBS and stained for p-AURAK (green), ARL13b (red), and γ-Tubulin (magenta). MEFs cultured in FBS are shown as a positive control for p-AURAK staining.

**Figure S3:**
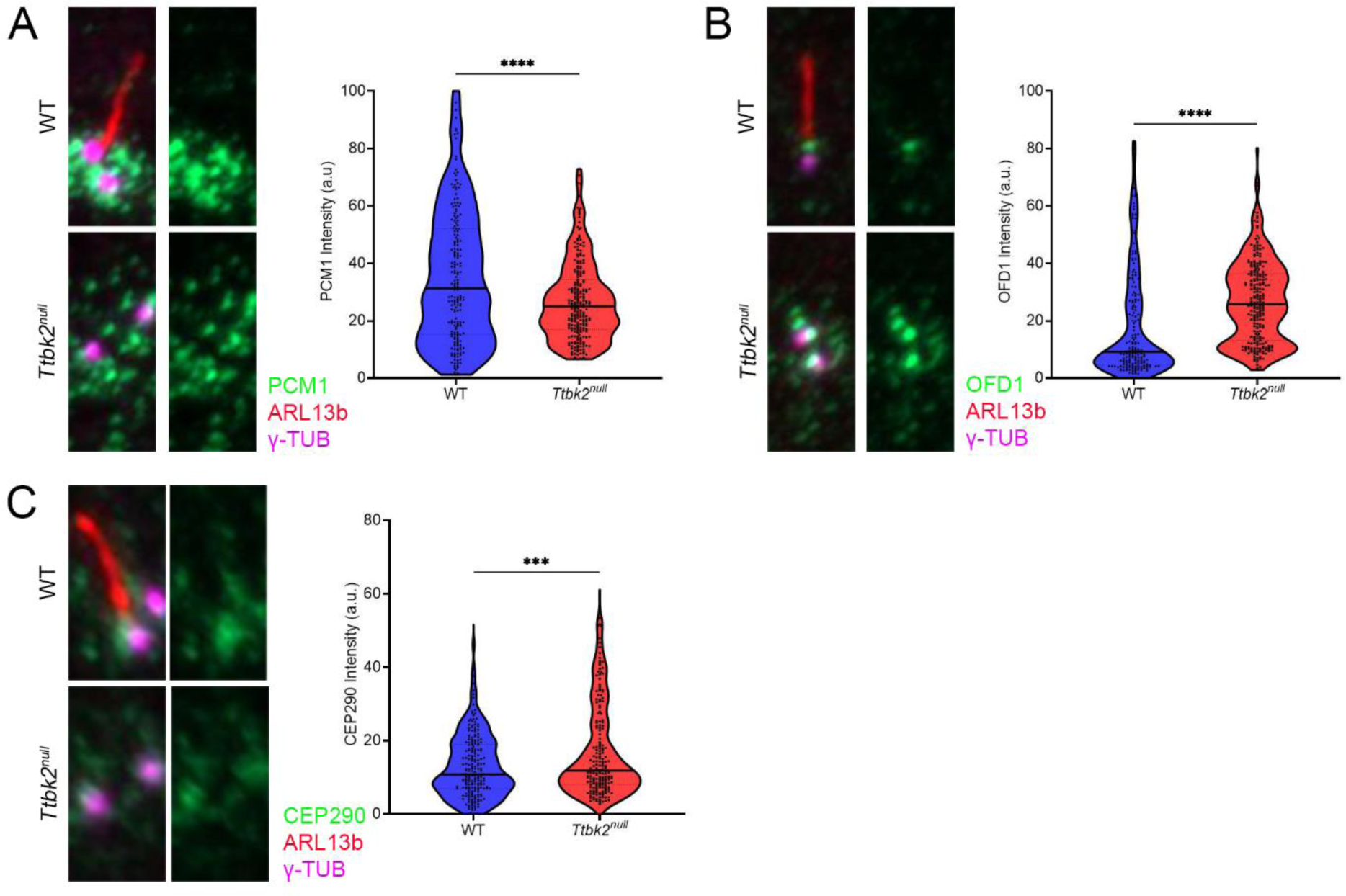
TTBK2 regulates the centriolar satellite composition during cilium initiation. A. Representative images of WT and *Ttbk2^null^* MEFs serum starved for 48hrs and stained for PCM1 (green), ARL13b (red), and γ-Tubulin (magenta). Graph depicts mean intensity of PCM1 1μm around basal bodies. Results are pooled from three independent experiments. Each dot represents a single measurement from a basal body (n>75). Statistical comparison was performed by a student’s unpaired t-test, **** denotes p<0.0001. B. Representative images of WT and *Ttbk2^null^* MEFs serum starved for 48hrs and stained for OFD1 (green), ARL13b (red), and γ-Tubulin (magenta). Graph depicts mean intensity of OFD1 1μm around basal bodies. Results are pooled from three independent experiments. Each dot represents a single measurement from a basal body (n>75). Statistical comparison was performed by a student’s unpaired t-test, **** denotes p<0.0001. C. Representative images of WT and *Ttbk2^null^* MEFs serum starved for 48hrs and stained for CEP290 (green), ARL13b (red), and γ-Tubulin (magenta). Graph depicts mean intensity of CEP290 1μm around basal bodies. Results are pooled from three independent experiments. Each dot represents a single measurement from a basal body (n>75). Statistical comparison was performed by a student’s unpaired t-test, *** denotes p<0.001.

**Figure S4.**
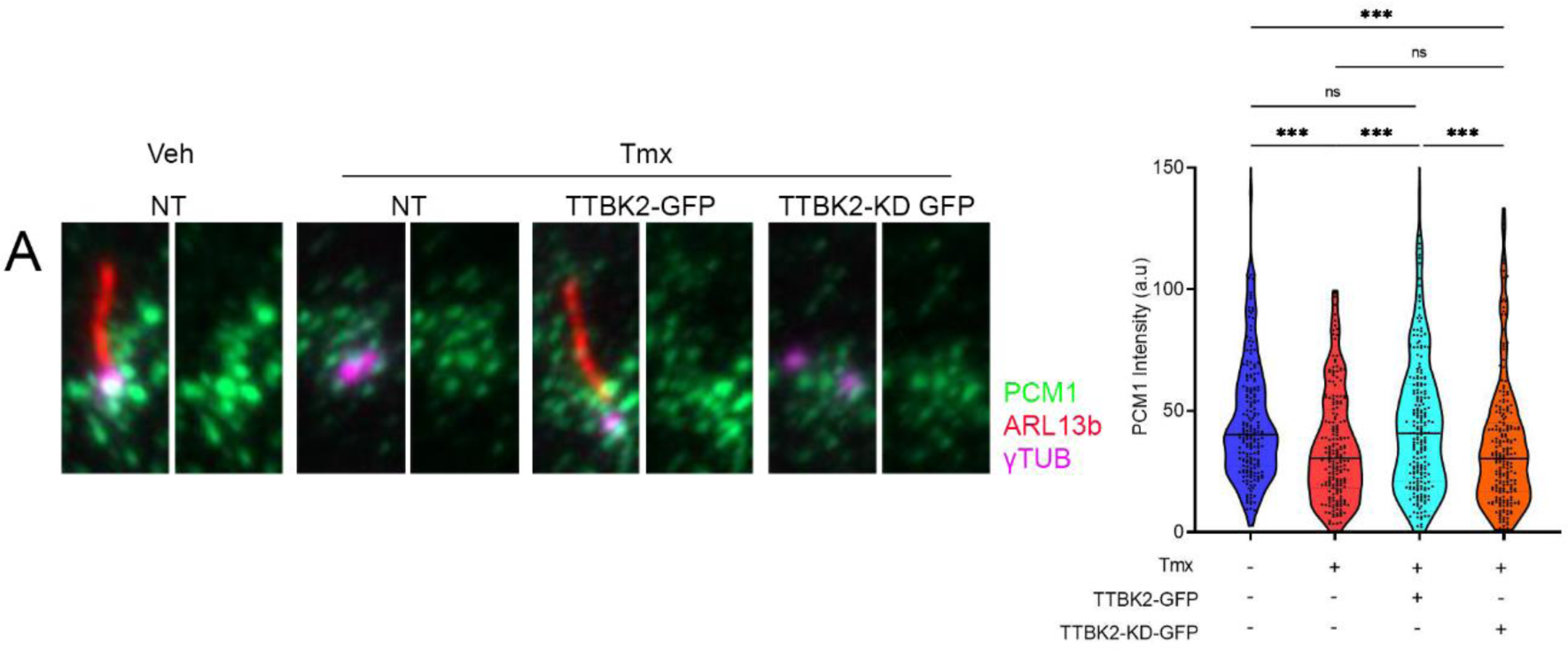
TTBK2’s kinase activity is required for centriolar satellite regulation. A. Representative images of *Ttbk2^cmut^* MEFs non-transfected (NT) or stably expressing TTBK2-GFP and TTBK2-KD-GFP treated with Veh or Tmx for 72 hours and stained for PCM1 (green), ARL13b (red), and γ-Tubulin (magenta). Graph depicts mean intensity of PCM1 1μm around basal bodies. Results are pooled from three independent experiments. Each dot represents a single measurement from a basal body (n>181). Statistical comparison was performed by one-way ANOVA with Tukey’s multiple comparisons test, ns denotes not significant, ** denotes p<0.01, *** denotes p<.001.

## References

1. Goetz SC, Anderson KV. The primary cilium: a signalling centre during vertebrate development. Nat Rev Genet. 2010 May;11(5):331–44.

2. Hilgendorf KI, Johnson CT, Jackson PK. The primary cilium as a cellular receiver: organizing ciliary GPCR signaling. Curr Opin Cell Biol. 2016 Apr;39:84–92.

3. Sánchez I, Dynlacht BD. Cilium assembly and disassembly. Nat Cell Biol. 2016 Jun 28;18(7):711–7.

4. Bowler M, Kong D, Sun S, Nanjundappa R, Evans L, Farmer V, et al. High-resolution characterization of centriole distal appendage morphology and dynamics by correlative STORM and electron microscopy. Nat Commun. 2019 Mar 1;10(1):993.

5. Tanos BE, Yang H-J, Soni R, Wang W-J, Macaluso FP, Asara JM, et al. Centriole distal appendages promote membrane docking, leading to cilia initiation. Genes Dev. 2013 Jan 15;27(2):163–8.

6. Singla V, Romaguera-Ros M, Garcia-Verdugo JM, Reiter JF. Ofd1, a human disease gene, regulates the length and distal structure of centrioles. Dev Cell. 2010 Mar 16;18(3):410–24.

7. Čajánek L, Nigg EA. Cep164 triggers ciliogenesis by recruiting Tau tubulin kinase 2 to the mother centriole. Proc Natl Acad Sci U S A. 2014 Jul 15;111(28):E2841–50.

8. Craft JM, Harris JA, Hyman S, Kner P, Lechtreck KF. Tubulin transport by IFT is upregulated during ciliary growth by a cilium-autonomous mechanism. J Cell Biol. 2015 Jan 19;208(2):223–37.

9. Nakayama K, Katoh Y. Ciliary protein trafficking mediated by IFT and BBSome complexes with the aid of kinesin-2 and dynein-2 motors. J Biochem. 2018 Mar 1;163(3):155–64.

10. Goetz SC, Liem KF Jr, Anderson KV. The spinocerebellar ataxia-associated gene Tau tubulin kinase 2 controls the initiation of ciliogenesis. Cell. 2012 Nov 9;151(4):847–58.

11. Bowie E, Norris R, Anderson KV, Goetz SC. Spinocerebellar ataxia type 11-associated alleles of Ttbk2 dominantly interfere with ciliogenesis and cilium stability. PLoS Genet. 2018 Dec 10;14(12):e1007844.

12. Bowie E, Goetz SC. TTBK2 and primary cilia are essential for the connectivity and survival of cerebellar Purkinje neurons. Elife [Internet]. 2020 Jan 14;9. Available from: http://dx.doi.org/10.7554/eLife.51166

13. Higginbotham H, Bielas S, Tanaka T, Gleeson JG. Transgenic mouse line with green-fluorescent protein-labeled Centrin 2 allows visualization of the centrosome in living cells. Transgenic Res. 2004 Apr;13(2):155–64.

14. Bangs FK, Schrode N, Hadjantonakis A-K, Anderson KV. Lineage specificity of primary cilia in the mouse embryo. Nat Cell Biol. 2015 Feb;17(2):113–22.

15. Loukil A, Barrington C, Goetz SC. A complex of distal appendage-associated kinases linked to human disease regulates ciliary trafficking and stability. Proc Natl Acad Sci U S A [Internet]. 2021 Apr 20;118(16). Available from: http://dx.doi.org/10.1073/pnas.2018740118

16. Mirvis M, Siemers KA, Nelson WJ, Stearns TP. Primary cilium loss in mammalian cells occurs predominantly by whole-cilium shedding. PLoS Biol. 2019 Jul 17;17(7):e3000381.

17. Wang G, Chen Q, Zhang X, Zhang B, Zhuo X, Liu J, et al. PCM1 recruits Plk1 to the pericentriolar matrix to promote primary cilia disassembly before mitotic entry. J Cell Sci. 2013 Mar 15;126(Pt 6):1355–65.

18. Miyamoto T, Hosoba K, Ochiai H, Royba E, Izumi H, Sakuma T, et al. The Microtubule-Depolymerizing Activity of a Mitotic Kinesin Protein KIF2A Drives Primary Cilia Disassembly Coupled with Cell Proliferation. Cell Rep. 2015 Feb 4;10(5):664–73.

19. Loskutov YV, Griffin CL, Marinak KM, Bobko A, Margaryan NV, Geldenhuys WJ, et al. LPA signaling is regulated through the primary cilium: a novel target in glioblastoma. Oncogene [Internet]. 2018 Jan 11; Available from: http://dx.doi.org/10.1038/s41388-017-0049-3

20. He K, Ling K, Hu J. The emerging role of tubulin posttranslational modifications in cilia and ciliopathies. Biophysics Reports. 2020 Aug 1;6(4):89–104.

21. He M, Subramanian R, Bangs F, Omelchenko T, Liem KF Jr, Kapoor TM, et al. The kinesin-4 protein Kif7 regulates mammalian Hedgehog signalling by organizing the cilium tip compartment. Nat Cell Biol. 2014 Jul;16(7):663–72.

22. Nager AR, Goldstein JS, Herranz-Pérez V, Portran D, Ye F, Garcia-Verdugo JM, et al. An Actin Network Dispatches Ciliary GPCRs into Extracellular Vesicles to Modulate Signaling. Cell. 2017 Jan 12;168(1–2):252–263.e14.

23. Mancini A, Sirabella D, Zhang W, Yamazaki H, Shirao T, Krauss RS. Regulation of myotube formation by the actin-binding factor drebrin. Skelet Muscle. 2011 Dec 8;1(1):36.

24. Shu S, Liu X, Korn ED. Blebbistatin and blebbistatin-inactivated myosin II inhibit myosin II-independent processes in *Dictyostelium*. Proc Natl Acad Sci U S A. 2005 Feb 1;102(5):1472–7.

25. Ran J, Yang Y, Li D, Liu M, Zhou J. Deacetylation of α-tubulin and cortactin is required for HDAC6 to trigger ciliary disassembly. Sci Rep. 2015 Aug 6;5:12917.

26. Pugacheva EN, Jablonski SA, Hartman TR, Henske EP, Golemis EA. HEF1-dependent Aurora A activation induces disassembly of the primary cilium. Cell. 2007 Jun 29;129(7):1351–63.

27. Kim J, Lee JE, Heynen-Genel S, Suyama E, Ono K, Lee K, et al. Functional genomic screen for modulators of ciliogenesis and cilium length. Nature. 2010 Apr 15;464(7291):1048–51.

28. Kohli P, Höhne M, Jüngst C, Bertsch S, Ebert LK, Schauss AC, et al. The ciliary membrane-associated proteome reveals actin-binding proteins as key components of cilia. EMBO Rep. 2017 Sep;18(9):1521–35.

29. Magistrati E, Maestrini G, Niño CA, Lince-Faria M, Beznoussenko G, Mironov A, et al. Myosin VI regulates ciliogenesis by promoting the turnover of the centrosomal/satellite protein OFD1. EMBO Rep. 2021 Dec 27;e54160.

30. Odabasi E, Gul S, Kavakli IH, Firat-Karalar EN. Centriolar satellites are required for efficient ciliogenesis and ciliary content regulation. EMBO Rep [Internet]. 2019 Apr 25; Available from: http://dx.doi.org/10.15252/embr.201947723

31. Tang Z, Lin MG, Stowe TR, Chen S, Zhu M, Stearns T, et al. Autophagy promotes primary ciliogenesis by removing OFD1 from centriolar satellites. Nature. 2013 Oct 10;502(7470):254–7.

32. Hori A, Toda T. Regulation of centriolar satellite integrity and its physiology. Cell Mol Life Sci. 2017 Jan;74(2):213–29.

33. Bouskila M, Esoof N, Gay L, Fang EH, Deak M, Begley MJ, et al. TTBK2 kinase substrate specificity and the impact of spinocerebellar-ataxia-causing mutations on expression, activity, localization and development. Biochem J. 2011 Jul 1;437(1):157–67.

34. Watanabe R, Wei L, Huang J. mTOR signaling, function, novel inhibitors, and therapeutic targets. J Nucl Med. 2011 Apr;52(4):497–500.

35. Melick CH, Jewell JL. Regulation of mTORC1 by Upstream Stimuli. Genes [Internet]. 2020 Aug 25;11(9). Available from: http://dx.doi.org/10.3390/genes11090989

36. Pigino G. Intraflagellar transport. Curr Biol. 2021 May 24;31(10):R530–6.

37. Ford MJ, Yeyati PL, Mali GR, Keighren MA, Waddell SH, Mjoseng HK, et al. A Cell/Cilia Cycle Biosensor for Single-Cell Kinetics Reveals Persistence of Cilia after G1/S Transition Is a General Property in Cells and Mice. Dev Cell. 2018 Nov 19;47(4):509–523.e5.

38. Liao J-C, Yang TT, Weng RR, Kuo C-T, Chang C-W. TTBK2: a tau protein kinase beyond tau phosphorylation. Biomed Res Int. 2015 Apr 9; 2015(4):575170.

39. Watanabe T, Kakeno M, Matsui T, Sugiyama I, Arimura N, Matsuzawa K, et al. TTBK2 with EB1/3 regulates microtubule dynamics in migrating cells through KIF2A phosphorylation. J Cell Biol. 2015 Aug 31;210(5):737–51.

40. Farina F, Ramkumar N, Brown L, Samandar Eweis D, Anstatt J, Waring T, et al. Local actin nucleation tunes centrosomal microtubule nucleation during passage through mitosis. EMBO J [Internet]. 2019 Jun 3;38(11). Available from: http://dx.doi.org/10.15252/embj.201899843

41. Inoue D, Obino D, Pineau J, Farina F, Gaillard J, Guerin C, et al. Actin filaments regulate microtubule growth at the centrosome. EMBO J [Internet]. 2019 Jun 3;38(11). Available from: http://dx.doi.org/10.15252/embj.201899630

42. Aydin ÖZ, Taflan SO, Gurkaslar C, Firat-Karalar EN. Acute inhibition of centriolar satellite function and positioning reveals their functions at the primary cilium. PLoS Biol. 2020 Jun;18(6):e3000679.

43. Seeley ES, Nachury MV. The perennial organelle: assembly and disassembly of the primary cilium. J Cell Sci. 2010 Feb 15;123(Pt 4):511–8.

44. Lopes CAM, Prosser SL, Romio L, Hirst RA, O’Callaghan C, Woolf AS, et al. Centriolar satellites are assembly points for proteins implicated in human ciliopathies, including oral-facial-digital syndrome 1. J Cell Sci. 2011 Feb 15;124(Pt 4):600–12.

45. Davenport JR, Watts AJ, Roper VC, Croyle MJ, van Groen T, Wyss JM, et al. Disruption of intraflagellar transport in adult mice leads to obesity and slow-onset cystic kidney disease. Curr Biol. 2007 Sep 18;17(18):1586–94.

46. Hibbard JVK, Vazquez N, Satija R, Wallingford JB. Protein turnover dynamics suggest a diffusion-to-capture mechanism for peri-basal body recruitment and retention of intraflagellar transport proteins. Mol Biol Cell. 2021 Jun 1;32(12):1171–80.

47. Lo C-H, Lin I-H, Yang TT, Huang Y-C, Tanos BE, Chou P-C, et al. Phosphorylation of CEP83 by TTBK2 is necessary for cilia initiation. J Cell Biol [Internet]. 2019 Aug 27; Available from: http://dx.doi.org/10.1083/jcb.201811142

48. Bernatik O, Pejskova P, Vyslouzil D, Hanakova K, Zdrahal Z, Cajanek L. Phosphorylation of multiple proteins involved in ciliogenesis by Tau Tubulin kinase 2. Mol Biol Cell. 2020 Mar 4;mbcE19060334.

